# Primary tumor associated macrophages activate programs of invasion and dormancy in disseminating tumor cells

**DOI:** 10.1101/2021.02.04.429798

**Authors:** Lucia Borriello, Anouchka Coste, Ved P. Sharma, George S. Karagiannis, Yu Lin, Yarong Wang, Xianjun Ye, Camille L. Duran, Xiaoming Chen, Erica Dalla, Deepak K. Singh, Maja H. Oktay, Julio A. Aguirre-Ghiso, John Condeelis, David Entenberg

**Author notes:** Corresponding Authors: John Condeelis, David Entenberg.

## Abstract

Metastases are initiated by disseminated tumor cells (DTCs) that depart from the primary tumor and colonize target organs. Growing evidence suggests that the microenvironment of the primary tumor lesion primes DTCs to display dormant or proliferative fates in target organs. However, the manner in which events taking place in the primary tumor influence DTC fate, sometimes long after dissemination, remains poorly understood. With the advent of a novel intravital imaging technique called the Window for High-Resolution Intravital Imaging of the Lung (WHRIL), we have, for the first time, been able to study the live lung longitudinally and follow the fate of individual DTCs that spontaneously disseminate from orthotopic breast tumors. We find, across several models, a high rate of success for tumor cells to complete the initial steps of the metastatic cascade in the secondary site, including retention of DTCs in the lung vasculature, speed of extravasation, and survival after extravasation. Importantly, initiation of metastatic growth was controlled primarily by a rate-limiting step that occurred post-extravasation and at the stage of the conversion of single DTCs from a dormant to a proliferative state. Detailed analysis of these events revealed that, even before dissemination, a subset of macrophages within the primary tumor induces, in tumor cells that are about to disseminate, the expression of proteins that regulate a pro- dissemination (Mena^INV^) and pro-dormancy (NR2F1) phenotype. Surprisingly, if cancer cells are intravenously injected, the rate limiting stages of Mena^INV^-associated extravasation, dormancy, and other parameters, are lost or altered in a way that impacts how DTCs progress through the metastatic cascade. Our work provides novel insight into how specific primary tumor microenvironments prime a subpopulation of cells for dissemination and dormancy. We also propose that dissecting mechanisms of metastasis, or testing anti-metastatic therapies, may yield results of limited application if derived from models that do not follow spontaneous dissemination.

**SIGNIFICANCE:** This study provides important insight into the contribution of primary tumor microenvironmental niches to cancer metastasis by identifying the manner in which these niches spawn subpopulations of DTCs that are primed for dissemination and dormancy in the secondary site. This study may provide novel targets that could be inhibited to prevent successful colonization of the secondary site and, hence, metastasis.

## INTRODUCTION

Metastasis causes approximately 90% of cancer-related mortality (1). It is the endpoint of a complex and dynamic cascade of steps in which tumor cells migrate within the primary tumor, intravasate, disseminate via the circulatory system, arrest at a secondary site, extravasate, survive, and finally re- initiate growth to form secondary tumors (2). Most therapeutic interventions targeting metastasis are designed only to attack actively dividing tumor cells (i.e. those in the last step of this cascade; active growth) and do not block any of the intermediate steps. Understanding the mechanisms that regulate the ability of disseminated tumor cells (DTCs) to complete all steps of metastasis will reveal additional targets for novel anti-metastatic therapeutics (3).

Since 1900, a number of studies have attempted to measure the fate of DTCs during the metastatic cascade using a variety of techniques. These include: histopathology of secondary sites such as the lung (4, 5); radioactive labeling and fate-mapping of DTCs in mice (6, 7); *ex vivo* visualization of DTCs in tissue explants (8); and *in vivo* visualization of DTCs in the Chick Chorioallantoic Membrane (CAM) (9, 10) and in secondary sites in mice such as the liver (11), brain (12), and lung (13). However, the results of these studies remain contradictory, alternatively identifying tumor cell survival in the circulation (7, 14), extravasation (15), and re-initiation of growth/initiation of dormancy (16, 17) as rate limiting steps.

Furthermore, the vast majority of these studies have relied on a technique called “experimental metastasis” (EM). EM is a process in which tumor cells are injected directly into the vasculature of mice (4) and is used in place of spontaneous metastasis (SM), where tumor cells in an orthotopic primary tumor stochastically progress through all of the steps of the metastatic cascade (4). EM assays assume that tumor cells injected as a bolus directly into the vasculature will have the same fate (and provide the same information on metastasis initiation) as do tumor cells which originate from within a primary tumor. However, a growing literature shows that the presence of a primary tumor can have a profound influence on metastatic outcome. For example, it has been determined that gene expression signatures in the primary tumor can provide information on the potential for metastatic relapse years after the primary tumor has been removed (18–20). Furthermore, specific primary tumor microenvironments, such as hypoxia, can prime those primary tumor cells that are destined to become DTCs to have divergent fates in target organs (21). Despite these advances, it is not known what influence the primary tumor has on the intermediate steps of the metastatic cascade in the secondary site.

In the past, we, and others, have used high-resolution intravital imaging (IVI) to investigate the process of intravasation and dissemination in the primary tumor (22, 23). To understand the influence the primary tumor has on spontaneously disseminating DTCs in the secondary site, we recently developed the implantable Window for High-Resolution Imaging of the Lung (WHRIL) (24). The WHRIL is the first method capable of providing a longitudinal view of the living lung at sub-cellular resolution over days to weeks. Here, we employed this tool to, for the first time, directly and longitudinally visualize the secondary site with single-cell resolution IVI throughout the process of spontaneous metastasis in order to directly measure the primary tumor’s influence on the initial steps of the metastatic cascade in the secondary site.

## RESULTS

### Real Time Observation of Tumor Cell Arrival to the Lung

We began our investigation by generating models of spontaneous metastasis. For these studies, we specifically chose to use orthotopic injections of cancer cell lines, and not transgenic animal models of cancer, so as to be able adhere closely to earlier studies of metastasis which all used cancer cell lines (7,15–17,25,26). Specifically, we developed orthotopic primary breast tumors in mice using E0771 cells labeled with green fluorescent protein (E0771-GFP) and grown in syngeneic C57/B6 mice (**Figure 1A, top**) or, to ensure that the observed biological findings were generalizable, using human tumor cells (MDA-MB-231) labeled with GFP (231-GFP) grown in nude immunocompromised mice. To observe DTCs in the lung, we implanted WHRILs into tumor-bearing mice, and 24 hrs later, imaged them with intravital imaging (IVI) using multiphoton microscopy (**Supplemental Movies 1 & 2**), as previously described (24). Using this method, we were able to visualize both intravascular and extravascular tumor cells (**Supplemental Figures 1A & B**) and to observe the arrival of new DTCs to the lung vasculature in real time (**Figure 1B, “Arrival”, top and bottom**).

**Figure 1:**
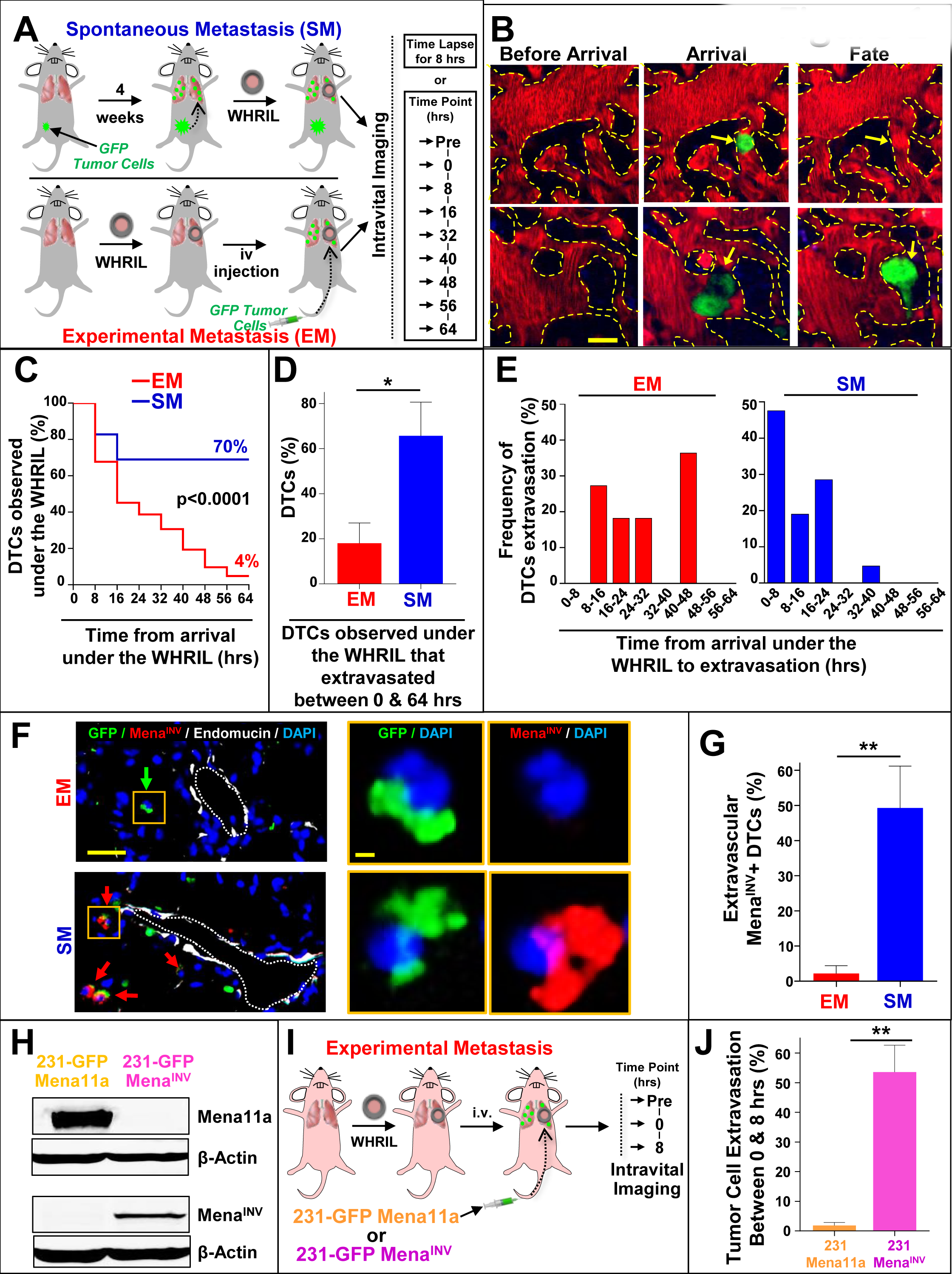
Tumor Cells that Spontaneously Disseminate from the Primary Tumor to the Lung have a Drastically Increased Metastatic Efficiency Compared to Intravenously Injected Tumor Cells. **A:** Outline of experimental design to track the fate of individual disseminated tumor cells using an Experimental Metastasis (EM) model (**top**) and a Spontaneous Metastasis (SM) model (**bottom**). In EM, the Window for High Resolution Intravital Imaging of the Lung (WHRIL) was surgically implanted in a tumor free mouse, and, 24 hrs after recovery from surgery, GFP labeled tumor cells were intravenously (iv) injected. In SM, GFP labeled tumor cells were injected into the mammary gland and tumors allowed to develop for ∼ 4 weeks after which the WHRIL was surgically implanted and the mouse allowed to recover for 24 hrs. For both models, intravital imaging consisted either of the acquisition of time-lapse images over an 8 hr period, or of the acquisition of single snap shot images every 8 hours using microcartography to return to the same field of view each time. In the case of snapshot imaging, an additional “Pre” time point is acquired before the others to visualize any preexisting tumor cells whose arrival time cannot be determined (SM model) or to visualize the empty lung vasculature just before iv injection (EM model). **B:** Serial imaging through the WHRIL allows tracking the fate of disseminated tumor cells. This is accomplished by imaging the lung to visualize the vasculature before the arrival of a tumor cell (**left**), and then again periodically to determine when a tumor cell first arrives (**middle, yellow arrow**). Continued periodic imaging then determines the fate of the tumor cell. This fate could be either recirculation or apoptosis (**right, top, yellow arrow**) or extravasation into the lung parenchyma (**right, bottom, yellow arrow**). Red = tdTomato labeled endothelial cells and 155 kDa Tetramethylrhodamine dextran labeled blood serum, Green = GFP labeled tumor cells, Blue = second harmonic generation. Scale bar = 15 μm. **C:** Kaplan-Meier curves showing the percentage of E0771-GFP tumor cells observed under the WHRIL at each 8 hr time point over a period of 64 hrs. EM: n = 62 tumor cells analyzed in 3 mice. SM: n = 29 tumor cells analyzed in 3 mice. **D:** Percentage of E0771-GFP EM and SM tumor cells observed under the WHRIL that extravasated between 0 and 64 hrs after arrival. EM: n = 11 tumor cells in 3 mice. SM: n = 22 tumor cells in 3 mice. Bar = mean. Error bars = ±SEM. * = p<0.05. **E:** Quantification of the time from arrival under the WHRIL to extravasation into the lung parenchyma for each E0771-GFP EM and SM tumor cell. **Left:** EM: n = 11 tumor cells in 4 mice. **Right:** SM: n = 22 tumor cells in 3 mice. **F: Left:** Representative immunofluorescence images of Mena^INV^ expression in extravascular E0771-GFP tumor cells in the lung of an EM model (**top**) and an SM model (**bottom**). Green arrow: Mena^INV^ negative tumor cell. Red arrows: Mena^INV^ positive tumor cells. Scale bar = 50 μm. **Right:** Zoomed in view of a disseminated tumor cell (yellow box) in both models. Green = GFP, Red = Mena^INV^, White = endomucin, Blue = DAPI. Scale bar = 15 μm. **G:** Quantification of extravascular Mena^INV^ positive disseminated tumor cells in the lung of each group from F. EM: n = 41 cells in 5 animals. SM: n = 89 cells in 7 animals. Bar = mean. Error bars = ±SEM. ** = p<0.01. **H:** Western blot of 231-GFP-Mena11a and 231-GFP-Mena^INV^ cells. β-Actin was used as a loading control. **I:** Outline of experimental design to determine the percentage of 231-GFP-Mena11a and 231-GFP- Mena^INV^ tumor cells able to extravasate between 0 and 8 hrs after iv-injection into nude mice. Intravital imaging of the lung vasculature through the WHRIL was performed before tumor cell injection (**Time point = “Pre”**), immediately after injection (**Time point = 0 hrs**), and then finally 8 hrs after injection (**Time point = 8 hrs**). **J:** Percentage of 231-GFP-Mena11a and 231-GFP-Mena^INV^ that extravasated between 0 and 8 hrs after iv injection. 231-GFP-Mena11a: n = 90 cells in 3 mice. 231-GFP-Mena^INV^: n = 88 tumor cells in 3 mice. Bar = mean. Error bars = ±SEM. ** = p<0.01.

Similar to observations made with vacuum-based imaging windows for the lung (13), we observed that tumor cells arriving to the lung vasculature completely fill the capillary lumen and exclude all blood serum (as indicated by IV-injected fluorescent dextran), suggesting that these cells are arrested due to physical restraint rather than active attachment. We did not find that tumor cells roll along the vasculature or otherwise migrate from their place of lodgment, as was previously observed in lung explants (8) or in zebrafish (27, 28). We never observed tumor cells to proliferate intravascularly, as reported by Al-Mehdi *et al.* (8). We also never observed CTC clusters traversing capillaries as was observed *in vitro* by Au *et al.* (29).

Since the progression of tumor cells through the metastatic cascade takes much longer than 8 hrs, we posited that the fate of disseminated tumor cells could be followed by imaging the vasculature once every 8 hrs, using *in vivo* microcartography (24) to relocate the same cells and microvasculature during each imaging session. To rule out the possibility that tumor cells may migrate in or out of the field of view between imaging sessions, we analyzed our time-lapse images to assess the motility of spontaneously metastasizing tumor cells before and after extravasation. As can be seen in **Supplemental Figures 1C Left & 1D Left**, we never observed tumor cells migrating beyond a single field of view. In fact, tumor cells were largely immobile. Based on our measurements, we projected that the fastest cells would take, at a minimum, 69 hrs to move from the center of the field of view to its boundary (256 µm). Thus, we next undertook evaluations of the long-term fate of disseminated tumor cells by capturing a single image of the vasculature once every eight hours.

### Spontaneously Metastasizing Tumor Cells are Retained in the Lung Significantly Longer and in Higher Numbers Compared to Intravenously Injected Tumor Cells

To track the fate of each newly arriving DTC as it progresses through the metastatic cascade, we prepared mice using the same procedure described above and then directly visualized the lung vasculature to record the presence of any previously disseminated tumor cells (**Figure 1A, Time Point “Pre”**), or the empty vasculature (**Figure 1B, Before Arrival**). The location of each preexisting tumor cell was recorded and these locations excluded from further analysis, as their precise arrival time was unknown. After 8 hrs, we again imaged the vasculature using the WHRIL, and recorded the locations of all newly arrived DTCs. These images constitute t=0 (**Figure 1A, Time Point 0 hrs**) with cells arriving sometime between 0 and 8 hrs after the start of the experiment.

We then imaged the lung vasculature every 8 hrs thereafter (**Figure 1A, Time Points 8 through 64 hrs**), using microcartography to predict to the same location for each time point. This location was verified by visual inspection of the unique architecture of the microvasculature. The fate of each tumor cell was determined as described in **Figure 1B**, which depicts the lung vasculature before the arrival of a tumor cell (**Figure 1B, “Before Arrival”, top and bottom**), the lodged tumor cell upon arrival (**Figure 1B, “Arrival”, top and bottom**), and finally the fate of the tumor cell as either death or recirculation within the vasculature (**Figure 1B, “Fate”, top**), or extravasation (**Figure 1B, “Fate”, bottom**).

With this methodology, we determined the arrival and subsequent disappearance or extravasation of tumor cells within 8 hr time frames. Unexpectedly, we found that, after an initial modest decline, most E0771-GFP tumor cells (>70%) were retained in the lung for the entire experimental period (**Figure 1C, blue curve**). Though, we observed a greater initial decline in the 231-GFP model, the number of tumor cells retained in the lung also persisted in this model, so that more than 35% remained after 64 hrs (**Supplemental Figure 2A, blue curve**). This was in stark contrast to previous reports where it was observed that DTCs would die off and be rapidly cleared from the tissue (7,25,30–32). To eliminate the use of the WHRIL as a contributing variable, we sought to repeat these experiments with the same experimental metastasis model used in the prior publications.

After repeating our time lapse imaging validation experiments to ensure that EM cells cannot migrate out of a field of view within 8 hours (**Supplemental Figures 1C Right & 1D Right**), we established a comparable method for tracking DTCs within the experimental metastasis model. We implanted WHRILs into mice and then 24 hrs later, we injected the same tumor cells into their tail veins (E0771-GFP in syngeneic C57/B6 or 231-GFP in nude mice). We then immediately began imaging the vasculature under the WHRIL. These images thus constitute t=0 for the EM model (**Figure 1A, bottom, Time Point 0 hrs**). We found that, as we had initially expected, E0771-GFP EM tumor cells were rapidly cleared from the lung, with a 50% drop within the first 16 hrs, and a steady decline thereafter, leaving only ∼4% after 64 hrs (**Figure 1C, red curve**). This represents a 10-fold increase in tumor cell retention of SM compared to EM tumor cells.

Consistently, we observed that the number of 231-GFP EM tumor cells also rapidly declined to 40% within the first 16 hrs, and then continued to decline steadily until less than 2% remained at 64 hrs post-injection (**Supplemental Figure 2A, red curve**). Although, the difference between EM and SM in this model was not as large as in the syngeneic model, the difference between the two models persisted and remained significant, confirming that disseminating tumor cells that originate in a primary tumor are retained longer and in higher numbers within the secondary site of the lung.

### Spontaneously Metastasizing Tumor Cells Extravasate More Quickly than Intravenously Injected Tumor Cells

It has been hypothesized that extravasation is a major rate-limiting step in metastasis because tumor cells are killed by the mechanical trauma that they are subjected to in the circulatory system (33, 34), and when tumor cells are able to extravasate quickly, they metastasize more efficiently (5). We therefore posited that the increased rate of retention of SM cells is due to a better ability to extravasate. To test this, we used the WHRIL to analyze the number of tumor cells that extravasated within the experimental time period (between 0 and 64 hrs). While only 19% of EM cells were able to extravasate, a significantly greater proportion of SM cells (64%) was able to extravasate during the same time period after arrival to the lung (**Figure 1D**). Interestingly, while we did observe a slight increase in the ability of SM cells to extravasate in the 231-GFP model, this was not statistically different from EM cells (**Supplemental Figure 2B**).

The observation that syngeneic SM cells extravasate more efficiently prompted us to evaluate how long individual tumor cells arriving in the lung take to cross the vascular endothelium. We therefore imaged SM and EM cells every 8 hrs after their arrival under the WHRIL and observed no EM cells to extravasate between 0 and 8 hrs after intravenous (iv) injection. The distribution of extravasation times was wide with many cells taking as long as 40-48 hrs to cross the endothelium (**Figure 1E, red bars**) with the average time of extravasation being 28 ± 4 hrs. In contrast, we found that a much larger proportion of SM cells was able to cross the endothelium quickly, with ∼50% doing so within 0-8 hrs from the time of first arrival to the lung vasculature (**Figure 1E, blue bars**). By 24 hrs, the vast majority of SM tumor cells had extravasated. Moreover, the average time of extravasation for SM cells was nearly half that for EM cells (12 ± 1.9 hrs, **Figure 1E**). Similar observations were made with 231-GFP cells, where only ∼30% of EM tumor cells extravasated within the 0-8 hr period while ∼80% of the SM cells extravasated in the same period (**Supplemental Figure 2C**). While the distribution of extravasation times was not as wide for the EM 231-GFP cells as was observed for the EM E0771-GFP cells, the difference in means persisted with EM cells extravasating on average at 11±1.0 hrs *vs.* 6±0.8 hrs for SM cells. These data demonstrate that disseminating tumor cells that originate in a primary tumor extravasate much more quickly than intravenously injected tumor cells.

### Expression of Mena^INV^ Confers Early Extravasation Competency to Disseminated Tumor Cells

Our previous studies (35–37) showed that expression of alternative splice variants of the actin regulatory protein, Mena, confer dramatically different metastatic phenotypes to tumor cells (38). Expression of the isoform, Mena11a, is associated with an epithelial, non-metastatic phenotype. Meanwhile Mena^INV^, an isoform not expressed in tumor cells cultured *in vitro*, but induced upon contact with macrophages, enhances tumor cell migration and transendothelial migration during intravasation within the primary tumor (35–37).

Therefore, we hypothesized that Mena^INV^ expression could play a similar role during extravasation at the secondary site by conferring to tumor cells the ability to cross the endothelium quickly. As a preliminary test, we took tissue sections from the lungs of the EM and SM models and stained them for GFP (to identify tumor cells), Mena^INV^, endomucin (to identify the vasculature), and DAPI (to identify all nuclei) (**Figure 1F**). We found a ∼25-fold increase (49% *vs.* 2%) in the percentage of SM tumor cells expressing Mena^INV^, compared to EM cells (**Figure 1G**). In the 231-GFP model, the percent of cells expressing Mena^INV^ was ∼4 times higher (48% *vs.* 13%) for SM compared to EM cells (**Supplemental Figure 2D&E**). This shows that Mena isoform expression in tumor cells retained in the lungs is significantly different between SM and EM cells.

To determine whether the expression of Mena^INV^ directly confers the ability to extravasate early (within 8 hr of arrival in the lung vasculature), we performed gain- and loss-of-function experiments using two genetically modified 231-GFP cell lines: one that overexpresses Mena^INV^ (231-GFP-Mena^INV^) and one that overexpresses Mena11a but does not express Mena^INV^ at all (231-GFP-Mena11a) (39) (**Figure 1H**). Using these two cell lines, we performed the experimental metastasis assay and tracked the number of tumor cells that had extravasated after 8 hr (**Figure 1I**). Over 50% of the 231-GFP-Mena^INV^ cells that arrived under the WHRIL extravasated within the first 8 hr post-injection while, lack of Mena^INV^ expression almost completely abrogated tumor cell extravasation within this time-period (**Figure 1J)**. This resulted in a nearly 30-fold increase in early extravasation of Mena^INV^ (54%) compared to Mena 11a (2%) expressing cells. These data show that the selective expression of Mena^INV^ is a requirement for early extravasation of DTCs into secondary sites.

### Spontaneously Metastasizing Tumor Cells Survive Significantly Longer at the Secondary Site Compared to Intravenously Injected Tumor Cells

It is generally accepted that after extravasation into the lungs, only a very small percentage of tumor cells survives (40). Given the differences between SM and EM observed thus far, we aimed to test whether the presence of the primary tumor influences tumor cell survival after extravasation in the lung. Thus, we imaged the lung vasculature in the SM and EM models, as described in **Figure 1A**, every eight hours for five days (120 hrs total) to track each individual tumor cell longitudinally. There were three possible outcomes for disseminated tumor cells: they died (**Figure 2A, left)**, evidenced by the presence of GFP+ cellular debris in the parenchyma, as was previously observed by Kienast et al. (12) (**Figure 2A, left, yellow arrow)**; they survived in the lung parenchyma as single cells (**Figure 2A, middle)**; or they grew into micro-metastases (**Figure 2A, right**).

**Figure 2:**
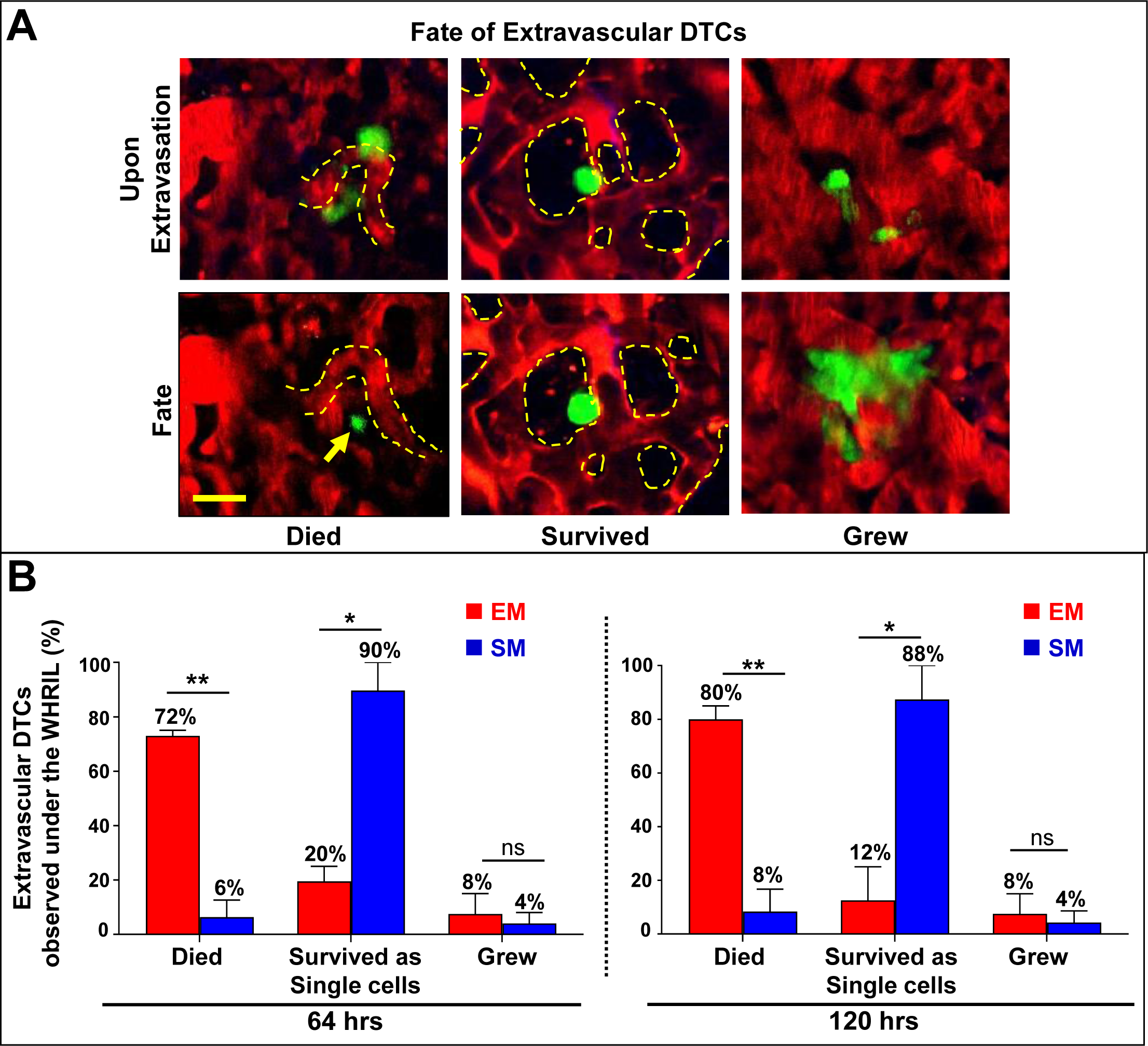
Spontaneously Metastasizing Tumor Cells Survive Significantly Longer at the Secondary Site Compared to Intravenously Injected Tumor Cells. **A:** Representative intravital microscopy images showing the possible fates of extravascular disseminated tumor cells in the lung parenchyma. **Top:** Images of disseminated tumor cells just after extravasation. **Bottom left:** Example of an extravascular tumor cell which has died, as evidenced by small extravascular apoptotic bodies (**yellow arrow**). **Bottom middle:** Example of an extravascular tumor cell that survived as a single and solitary tumor cell over time. **Bottom right:** Example of an extravascular tumor cell that began to divide and grow into a micro-metastasis. Red = tdTomato labeled endothelial cells and 155 kDa Tetramethylrhodamine dextran labeled blood serum, Green = GFP labeled tumor cells. Yellow dashed lines delineate blood vessel boundaries. Scale bar = 15 μm. **B:** Percentage of extravascular E0771-GFP disseminated tumor cells that died, survived, or grew after extravasation in EM and SM models 64 hrs (**Left**) and 120 hrs (**Right**) after arrival to the lung vasculature. EM: n = 11 tumor cells in 4 mice. SM: n = 22 tumor cells in 3 mice. Bar = mean. Error bars = ±SEM. * = p<0.05. ** = p<0.01. ns = not significant.

We found that a large majority (72%) of EM cells died shortly after extravasation, with a 28% survival rate at 64 hrs. Of these, 20% remained as single cells at 64 hrs post-injection and 8% began to form micro-metastasis **(Figure 2B left, red bars)**. When followed out to five days, the number of EM cells that died grew slightly to 80% while the number that remained as single cells nearly halved (12%) (**Figure 2B right, red bars**). The 8% of cells that had started to grow remained constant over this longer period. In contrast, by 64 hrs after extravasation, the vast majority of SM cells (90%) survived as single and solitary tumor cells and only a small percentage (6%) died. Of the surviving cells, only a small subset formed micro-metastasis (4%) during this time period (**Figure 2B left, blue bars**). The number of SM cells that died increased only slightly (from 6 to 8%) by five days, due solely to the death of single, non-dividing cells (going from 90 to 88%). The number of growing SM cells was unchanged between the two time points (**Figure 2B right, blue bars**). Similar observations were made with 231-GFP cells (**Supplemental Figure 2F**). These data demonstrate that disseminated tumor cells originating in a primary tumor have a drastically increased ability to extravasate, then seed and successfully survive in the lung parenchyma compared to intravenously injected tumor cells.

### Spontaneously Metastasizing Tumor Cells Enter Dormancy More Frequently Than Intravenously Injected Tumor Cells

The observation that the vast majority of SM cells survive as single and solitary cells, without dying or proliferating during the first five days of their residency in the lung, suggested that they may have become dormant. This is consistent with earlier studies using EM to disseminate tumor cells in the CAM (41, 42), liver (42), and lung (17). However, in these studies, the dormant state of the tumor cells was only determined using rudimentary assessments (absence of division, Ki-67 expression). This prompted us to assess if the solitary DTCs we found in the SM model are also in a dormant state using more recently discovered markers. One of the best molecular markers of dormancy is the orphan nuclear receptor NR2F1 (41, 42), a member of the of the retinoic acid receptor family (42) that has been shown to be a marker for cancer dormancy in pre-clinical models and cancer patients, as well to be an independent prognostic indicator for time-to-distant-recurrence in breast cancer patients (41). Furthermore, NR2F1 has been shown to regulate tumor dormancy in different mouse models, including breast cancer (20,21,42,43).

In tissue and blood samples from both metastasis models, we could find single tumor cells (GFP+) that expressed nuclear NR2F1 (**Figure 3A**). We found that in lung tissues, the frequency of disseminated tumor cells expressing nuclear NR2F1 was upregulated ∼3-fold in SM cells when compared to EM cells (53% *vs.* 18%) (**Figure 3B, “Lung” columns**), suggesting that a significantly greater proportion of spontaneously disseminated tumor cells enter dormancy. We also observed numerous instances of NR2F1-positive tumor cells located inside the lung vasculature (**Supplemental Figure 3**). This raised the question of whether SM tumor cell dormancy was initiated after arrival to the lung vasculature, or if it occurred within the primary tumor, as we previously showed happens under hypoxic conditions (21). We therefore quantified NRF21+ cells in primary tumor tissues and circulating tumor cells (CTCs) from the SM model (**Figure 3A, Supplemental Figure 4A**). Only a very small percentage (∼3%) of primary tumor cells were positive for NR2F1 (**Figure 3B, “Primary Tumor” bar**), consistent with data from head and neck squamous cell carcinoma (HNSCC) PDX models (21) and the mouse mammary tumor virus-polyoma virus middle T antigen (MMTV-PyMT) model (21). Despite there being only a small number of NR2F1-positive primary tumor cells, we observed that ∼50% of CTCs were NR2F1-positive (**Figure 3B, “CTCs” bar**), suggesting that tumor cells are programmed for dormancy either before they intravasate (since acquisition can occur in hypoxic microenvironments (21)) or during intravasation. This represents a very significant enrichment (∼17-fold) upon intravasation. Given the short residence time of the cells in the vasculature as CTCs, it is unlikely this enrichment is due to the death of non-dormant cells within circulation. This high level of NR2F1 DTC positivity was also found inside of the lung tissue (**Figure 3B, left “Lung” bar**). Importantly, we did not observe a difference between the percentage of NR2F1-positive cells present *in vitro*, before injection, and the fraction of EM cells observed in the lung three days post injection (**Figure 3B, right, “in vitro” & “Lung” bars**), suggesting that expression of NR2F1 is not influenced by passage through the blood. Similar observations were made with 231-GFP cells (**Supplemental Figure 4B**), though the percentage of NR2F1 cells within the SM lung was significantly lower than in the CTCs. Overall, these data show that a larger proportion of tumor cells that originate in a primary tumor and arrive to the lungs are dormant compared to intravenously injected tumor cells.

**Figure 3:**
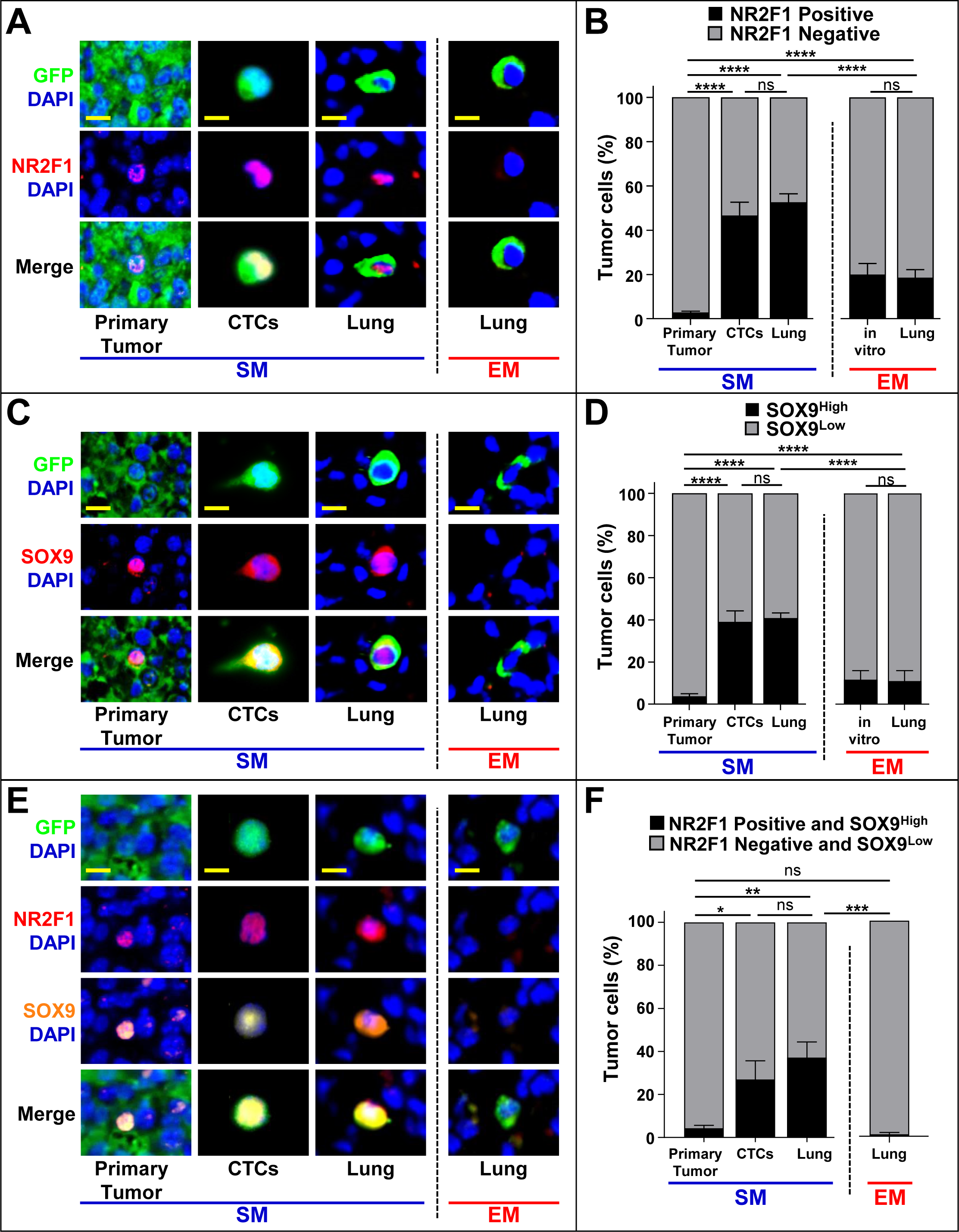
Spontaneously Metastasizing Tumor Cells Are More Frequently Positive for Dormancy and Stem-like Markers Compared to Intravenously Injected Tumor Cells. **A:** Representative immunofluorescence images of NR2F1 expression in primary tumors, circulating tumor cells (CTCs), and disseminated tumor cells (Lung) from an E0771-GFP SM model (**Left**) and in disseminated tumor cells (Lung) from an EM model (**right**). Green = GFP, Red = NR2F1, Blue = DAPI. Scale bar for Primary Tumor = 50 μm. Scale bar for CTCs and Lung = 15 μm. **B:** Percentage of NR2F1-positive and negative tumor cells in each group in **A**. Primary Tumor: n = 113 fields of view (65x65 µm^2^) in 8 animals; CTCs: n = 528 cells in 8 animals; SM Lung: n = 237 cells in 12 animals; EM Lung: n = 199 cells in 8 animals. In vitro: n= 463 cells in 5 independent experiments. Bar = mean. Error bars = ±SEM. **** = p<0.0001. ns = not significant. **C:** Representative immunofluorescence images of SOX9 expression in primary tumors, circulating tumor cells (CTCs), and disseminated tumor cells (Lung) from an E0771-GFP SM model (**Left**) and in disseminated tumor cells (Lung) from an EM model (**right**). Green = GFP, Red = SOX9, Blue = DAPI. Scale bar for Primary Tumor = 50 μm. Scale bar for CTCs and Lung = 15 μm. **D:** Percentage of SOX9^High^ tumor cells from each group in **C**. Primary Tumor: n = 150 fields of view (65x65 µm^2^) in 8 animals; CTCs: n = 558 cells in 5 animals, SM Lung: n = 341 cells in 11 animals; EM Lung: n = 182 cells in 8 animals. In vitro: n = 298 cells in 3 independent experiments. Bar = mean. Error bars = ±SEM. **** = p<0.0001. ns = not significant. **E:** Representative images of triple immunofluorescence staining for GFP, NR2F1, and SOX9 expression in primary tumors, circulating tumor cells (CTCs), and disseminated tumor cells (Lung) from an E0771- GFP SM model (**Left**) and in disseminated tumor cells (Lung) from an EM model (**Right**). Green = GFP; Red = NR2F1; Orange = SOX9; Blue = DAPI. Scale bar for Primary Tumor = 50 μm. Scale bar for CTCs and Lung = 15 μm. **F:** Percentage of double positive tumor cells NR2F1-positive SOX9^High^ from each group in **E**. Primary Tumor: n = 93 fields of view (65x65 µm^2^) in 7 animals; CTCs: n = 265 cells in 6 animals; SM Lung: n = 104 cells in 9 animals; EM Lung: n = 75 cells in 7 animals. Bar = mean. Error bars = ±SEM. * = p<0.05. ** = p<0.01. *** = p<0.001. ns = not significant.

### Spontaneously Metastasizing Tumor Cells Are More Frequently Positive for Both Dormancy and Stem-like Markers Compared to Intravenously Injected Tumor Cells

Recently, it was shown that NR2F1 in tumor cells coordinates the expression of other genes (e.g. SOX9, SOX2, and NANOG (44)) that are found in self-renewing embryonic stem cells and that can themselves coordinate quiescence (42). In particular, it was discovered that NR2F1 binds directly to the promoter of SOX9 to regulate SOX9 expression in tumor cells, resulting in dormancy and growth arrest (45). Furthermore, it was observed that SOX2 mRNA is significantly upregulated in dormant tumor cells (42). Based on these observations, we hypothesized that tumor cells originating in a primary tumor may take on a more stem-like phenotype compared to intravenously injected tumor cells as part of the dormancy program that is induced in the primary tumor.

To address this, lung tissue sections from EM and SM models, primary tumor tissues, and CTCs from the SM model, were stained for SOX9 (**Figure 3C**). The expression of SOX9 in SM cells in the lung was ∼4-fold higher when compared to EM cells (40% *vs.* 11%, **Figure 3D**). As with NR2F1, we found only a small population (∼4%) of SOX9^High^ cells in the primary tumor (**Figure 3D, “Primary Tumor**” bar), but a dramatic enrichment of SOX9^High^ CTCs (∼40%, a 10-fold increase, **Figure 3D, “CTCs**” bar), suggesting that tumor cells are programmed for stemness in the primary tumor, before, or during intravasation. Again, we did not observe a difference between the percentage of SOX9^High^ cells *in vitro* and the fraction of EM cells in the lung (**Figure 3B, right, “in vitro” & “Lung” bars**), suggesting that induction of a stem-like program does not occur in the blood. Similar observations were made with 231- GFP cells (**Supplemental Figure 4C&D**).

Given the mechanistic link between NR2F1 and SOX9 expression (42), we determined whether disseminated tumor cells co-express NR2F1 and SOX9, which would result in both quiescence and self- renewal (a stem-like program such as exists in adult quiescent stem cells). Lung tissue from EM and SM models, primary tumor tissues, and CTCs from the SM model were stained for GFP, NR2F1, SOX9, and DAPI (**Figure 3E & Supplemental Figure 4E**). In the primary tumor, we observed only a small population (∼4%) that co-expressed NR2F1 and SOX9. However, SM cells were enriched for double positive CTCs and DTCs in the lung compared EM cells (37% *vs.* 1%, **Figure 3F**). Similar observations were made with 231-GFP cells (**Supplemental Figure 4F**). Overall, these data show that SM tumor cells become progressively more enriched in dormancy and stem-like phenotypes as they disseminate from the primary tumor to the lung.

### Dormant Tumor Cells Are Preferentially Associated with TMEM Doorways in the Primary Tumor

Given the dramatic increase in dormancy markers as tumor cells move from the primary tumor into the vasculature, we hypothesized that the dormancy program may be activated near to, or at, intravasation sites. Our prior work (46–50) has shown that tumor cells intravasate through cellular doorways in the vasculature called tumor microenvironment of metastasis (TMEM) doorways. This stable triple cell complex is composed of a Mena^High^ tumor cell, a Tie2^High^ macrophage, and a blood vessel endothelial cell, all in direct physical contact. We have shown that TMEM are the sole sites of breast cancer cell intravasation (51) and we have clinically validated the density of TMEM doorways as a prognostic marker for metastatic recurrence in breast cancer patients (50). Thus, programming for dormancy might be induced as migratory tumor cells approach, interact with, or reside in the vicinity of TMEM doorways.

Serial sections of primary breast tumor tissues were stained for TMEM doorways using a triple immunohistochemical stain (48, 50) (**Figure 4A, left and insets**) or for GFP (to identify tumor cells), NR2F1, and DAPI. Using digital whole slide scanning and tissue alignment software (see **Methods**), we were able to co-register the two slides down to the single-cell level and measure the relative distance from each NR2F1-positive tumor cell to the nearest TMEM doorway. Analysis of these distances revealed an ∼2.5-fold enrichment of NR2F1-positive tumor cells near TMEM doorways (0-80 µm), compared to tumor cells farther away (160-200 µm) (**Figure 4B**). Interestingly, NR2F1-positive tumor cell enrichment was specifically associated with TMEM doorways, as we did not find any enrichment around blood vessels lacking these structures (**Figure 4C**). These data indicate that tumor cells within the primary tumor, and in close proximity to TMEM doorways, upregulate dormancy.

**Figure 4:**
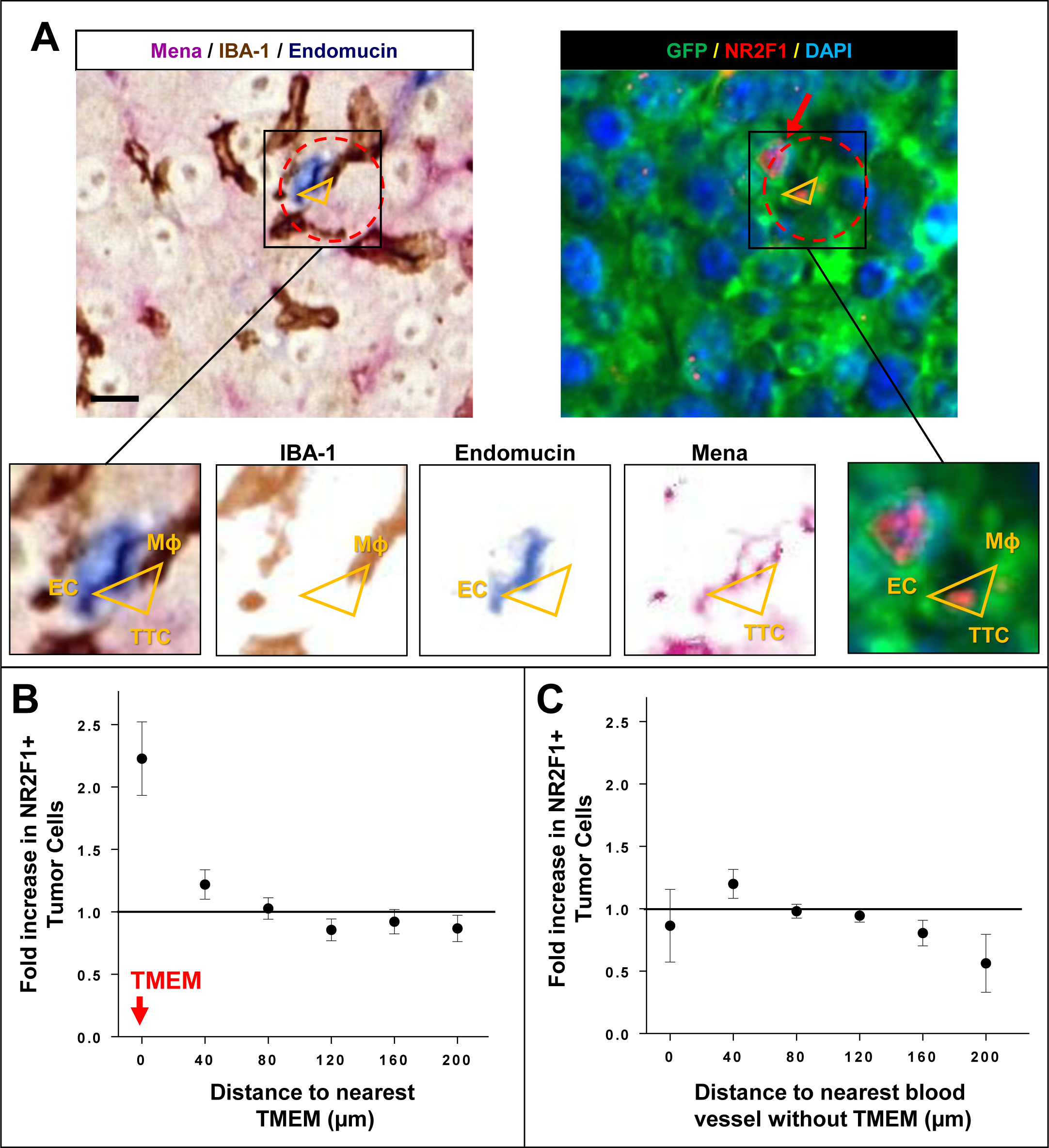
Dormant Tumor Cells Are Preferentially Associated with TMEM Doorways in the Primary Tumor. **A: Left:** Representative image of triple immunohistochemical stain in E0771-GFP primary tumor for the cells composing TMEM positioned at vertices of yellow triangle: Mena expressing tumor cells = pink; IBA- 1 expressing macrophages = brown; endomucin expressing endothelial cells = blue. Red dashed circle encompasses the perimeter of TMEM doorway. Scale bar = 60 µm. Insets are zoom-in of boxed region (**first panel on left**). Other insets show color deconvolutions for each of the stains (**Mena, Endomucin, IBA-1**). TTC = TMEM Tumor Cell. EC = Endothelial Cell. Mϕ = Macrophage. **Right:** Sequential slide of tissue in **A** immunofluorescently stained for NR2F1 expressing tumor cells: GFP = green; NR2F1 = red; DAPI = blue. Red dashed circle encompasses the perimeter of TMEM doorway. Vertices of the orange triangles point to each constitutive cell. Red arrow points to NR2F1-positive cells. **B:** Quantification showing frequency of distances between NR2F1^+^ tumor cells to the nearest TMEM in the primary tumor. Data is normalized to the frequency of distances between all DAPI^+^ nuclei to the nearest TMEM. Bar = mean. Error bars = ±SEM. n = ten 1-3 mm^2^ regions of interest area in 4 mice. **C:** Quantification showing frequency of distances between NR2F1^+^ tumor cells to the nearest blood vessel lacking TMEM in the primary tumor. Data is normalized to the frequency of distances between all DAPI^+^ nuclei to the nearest blood vessel lacking TMEM. Bar = mean. Error bars = ±SEM. n = nine 1-3 mm^2^ regions of interest area in 4 mice.

### Macrophages Regulate Dormancy in Disseminating Tumor Cells

To determine if tumor cell-macrophage interactions around TMEM induce tumor cell NR2F1- positivity, we fixed primary tumor tissues and studied the spatial distribution of dormant tumor cells relative to macrophages (**Figure 5A**). We observed an ∼2-fold enrichment of NR2F1-positive tumor cells in close proximity (0-40 µm) to macrophages in primary tumors. We also showed that NR2F1 expression is significantly increased in tumor cells co-cultured with macrophages compared to tumor cells cultured alone (48% *vs.* 11%), or co-cultured with endothelial cells (16% *vs.* 11%) (**Figure 5C&D**). When co-cultured cells were separated by a porous membrane, we observed a similar increase in tumor cell NR2F1 expression in the presence of macrophages, indicating that soluble factors are responsible for the induction of NR2F1 (**Supplemental Figure 5A&B).**

**Figure 5:**
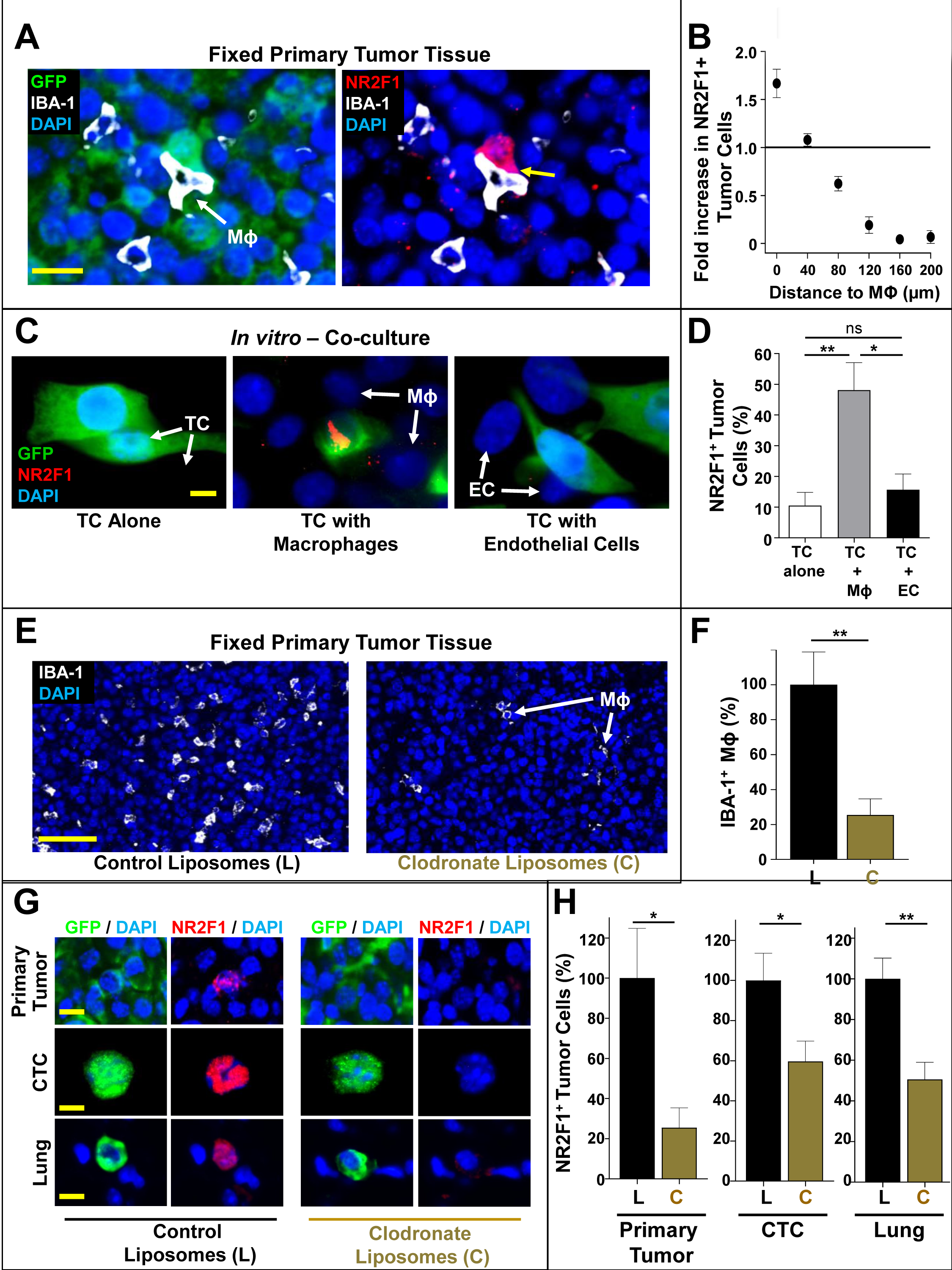
Macrophages Regulate Dormancy in Disseminating Tumor cells. **A:** Representative image of triple immunofluorescently stained in E0771-GFP primary tumor tissue for tumor cells, macrophages and NR2F1. Tumor cell GFP = green; NR2F1 = red; IBA-1 = white; DAPI = blue. Yellow arrow shows the contact between an NR2F1-positive tumor cell and a macrophage. Mϕ = Macrophage. Scale bar = 50 μm. **B:** Quantification showing frequency of distances between NR2F1^+^ tumor cells to the nearest macrophage in the primary tumor. Data is normalized to the frequency of distances between all DAPI^+^ nuclei to the nearest TMEM. Bar = mean. Error bars = ±SEM. n = 34 fields of view (551x316 µm^2^) in 4 animals. **C:** Representative immunofluorescence images of NR2F1 expression in E0771-GFP tumor cells cultured alone, in direct contact with BAC1.2F5 macrophages, or in direct contact with HUVEC endothelial cells. White arrows show macrophages or endothelial cells in direct contact with a tumor cell. GFP = green; NR2F1 = red; DAPI = blue. TC = Tumor Cell. Mϕ = Macrophage. EC = Endothelial Cell. Scale bar = 15 μm. **D:** Percentage of NR2F1-positive tumor cells from each group in **C**. TC alone: n = 637 cells in 9 independent experiments; TC + Mϕ; n = 195 cells in 6 independent experiments, TC + EC = n = 359 cells in 4 independent experiments. Bar = mean. Error bars = ±SEM. * = p<0.05. ** = p<0.01; ns = not significant. TC = Tumor Cell. Mϕ = macrophage. EC = Endothelial Cell. **E:** Representative immunofluorescence images of E0771-GFP primary tumor tissues treated for 6 days with either control liposomes or clodronate liposomes and stained for macrophages: IBA-1 = White; DAPI = Blue. Scale bar for Primary Tumor =100 μm. Mϕ = Macrophage. **F:** Percentage of IBA1 positive macrophages in 10 fields of view (1088x629 μm) in each group from **E**. Control Liposomes: n = 60 HPFs in 6 animals. Clodronate liposomes: n = 50 HPFs in 5 animals. ** = p<0.01. Bar = mean. Error bars = ±SEM. **G:** Representative immunofluorescence images of NR2F1 expression in primary tumors, circulating tumor cells (CTCs), and disseminated tumor cells (Lung) from an E0771-GFP SM model treated with control liposomes (**Left**) or with clodronate liposomes (**Right**). Green = GFP, Red = NR2F1, Blue = DAPI. Scale bar for Primary Tumor = 50 μm. Scale bar for CTCs and Lung = 15 μm. **H:** Percentage of NR2F1-positive tumor cells in each group from **G**. Control Liposomes: Primary Tumor: 119 fields of view (65x65 µm^2^) in 8 animals; CTCs: n = 139 cells in 5 animals; Lung: n = 166 cells in 7 animals. Clodronate Liposomes: Primary Tumor: n = 79 fields of view (65x65 µm^2^) in 6 animals, CTCs: n = 293 cells in 6 animals; Lung: n = 190 cells in 7 animals. * = p<0.05. ** = p<0.01.

Finally, we treated tumor-bearing mice (SM model) with control liposomes or clodronate liposomes to systematically deplete them of macrophages (52). Clodronate treatment led to significant macrophage depletion (∼40%) in primary tumor tissues (**Figure 5E&F, Supplemental Figure 5C&D**). We then determined the percentage of NR2F1-positive tumor cells in CTCs and lung tissues from an E0771-GFP SM model (**Figure 5G**). We found that, while the overall number of disseminated tumor cells is reduced in clodronate liposome treated animals (36,47,53), of the cells that do disseminate, there is a significant reduction in the fraction that are positive for NR2F1 in CTCs and in disseminated tumor cells found in the lungs, as well as in the primary tumor, compared to control animals (**Figure 5H**). Similar observations were made with 231-GFP cells (**Supplemental Figure 5E&F**). We therefore conclude that tumor associated macrophages play an important role in activating dormancy in disseminating tumor cells.

## DISCUSSION

Over the past decades, several studies have attempted to measure the fate of DTCs in secondary sites. These studies have inconsistently identified which steps are rate limiting in the metastatic cascade. In fact, almost all steps of metastasis have been identified, including intravasation, tumor cell survival in the circulation (7, 14), extravasation (15), and re-initiation of growth/initiation of dormancy (16, 17).

Studying the fate of individual DTCs in an intact organ such as the lung has been a major challenge for metastasis research because, until recently, it has not been possible to follow individual DTCs in the lung over time. Imaging techniques capable of viewing live lung tissue more than once (e.g. MRI, PET, CT) suffer from low resolution and are unable to visualize individual cells, while high resolution techniques only work on excised tissues and are thus limited to single time-point snapshots.

The result of these limitations is that it has been impossible to determine when spontaneously disseminated tumor cells had arrived to the organ, and for how long they had resided there. To overcome this limitation many studies have relied on the experimental metastasis (EM) assay where tumor cells are injected directly into the vasculature.

As revealed in our work, a major caveat of the EM assay is the implicit assumption that the processes of education that disseminating tumor cells undergo within the primary lesion are of marginal importance for DTC fate, and that cancer cells injected as a bolus directly into the vasculature are identical in their ability to progress through the metastatic cascade.

However, it is becoming increasingly clear that the primary tumor plays an important role in determining DTC fate. For example, it is possible to find gene signatures within the primary tumor that indicate whether tumor cells are likely to disseminate (54), and if they are likely to grow into metastases (55), even long after dissemination (19, 20). The primary tumor can also create systemic effects, preparing pre-metastatic niches in secondary sites (56) or influencing the reaction of the immune system to disseminated tumor cells (57). In addition to these effects, we recently determined that intratumoral microenvironments can activate programs of dormancy in DTCs (21) which may be the mechanism for therapy evasion and late recurrence in patients.

This is in accord with our observation that tumor cells that spontaneously disseminate from primary tumors remain in the lung for extended periods of time compared to those that are injected directly into the vasculature, suggesting that the primary tumor plays a protective role for these cells.

It was previously proposed that the harsh conditions of the circulatory system lead to tumor cell destruction (33), and that metastatic seeding is more likely to occur in areas with low perfusion (27). Consistent with this, we found that the survival advantage of spontaneously metastasizing (SM) cells over injected cells may be connected to their ability to extravasate into the lung parenchyma faster, resulting in a decreased time-from-arrival-to-extravasation and a shorter exposure to the circulatory system. Furthermore, our observation that a greater percentage of SM cells was able to extravasate in a syngeneic mouse model compared to an immunocompromised mouse model could be due to the immune system. While it not been shown that B and T cells influence DTC clearance from the lung vasculature, it has been shown that NK cells play a critical role in preventing tumor cell retention in the lung (58) (59) (5). Several studies demonstrated that the activity of NK cells is higher in nude mice compared to C57B6 mice (60) (61). It was furthermore shown that there is a difference in NK cell reactivity between nude and C57B6 mice with nude mice having much higher levels (60,62,63). Therefore, we argue that the observed increased clearance in the MD-MBA-231-eGFP SM model is likely due to a higher activity of NK cells in nude mice compared to C57B6 mice. This is an avenue that requires further investigation. Additionally, clearance of EM tumor cells from the vasculature was not a result of an immune reaction to GFP because an immune reaction would be expected to affect both EM and SM models equally, and it has been previously shown that GFP produces no detectable *in vivo* immune responses in C57B6 mice (64). Taken together, these observations indicate that neither destruction in the circulation nor extravasation are major limiting steps for disseminating tumor cells originating in a primary tumor.

Through gain- and loss-of-function experiments, we showed that Mena^INV^, an isoform of the actin regulatory protein Mena involved in cell motility and chemotaxis (38), is one important factor in the decreased time-to-extravasation of SM cells compared to EM cells. We previously showed that expression of Mena^INV^ drives invadopodium assembly and function (35), and is required for trans- endothelial migration within the primary tumor (65), and that the expression of Mena^INV^ persists in the primary tumor, CTCs, and DTCs in the lung (66). Our current work thus indicates that Mena^INV^ is a common molecular pathway, important for many of the steps of metastasis including invasion, intravasation, and now extravasation (**Figure 6**). While further studies will be required to elucidate the exact mechanism of Mena^INV^ induction, our prior work indicates that induction involves Notch mediated signaling resulting from macrophage-tumor cell contact in the primary tumor (35, 67).

**Figure 6:**
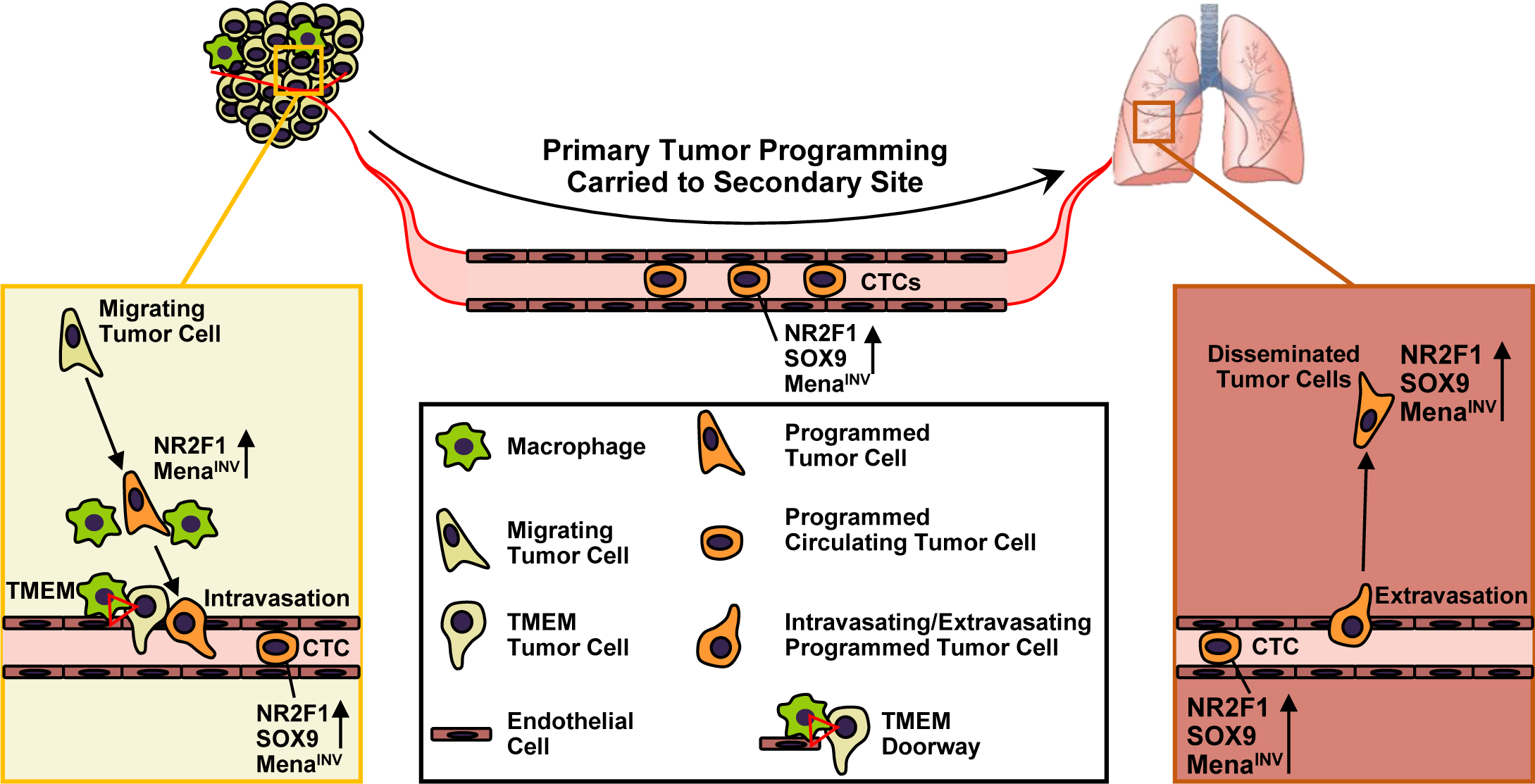
Model illustrating how the presence of a primary tumor programs disseminated tumor cells for stemness and dormancy at the secondary site. **Left Panel:** Within the primary tumor, migrating tumor cells are attracted to blood vessels. As they approach TMEM doorways (red triangle) on the vasculature, these tumor cells interact with macrophages and programs of dormancy (NR2F1) and invasion (Mena^INV^) are activated. Dormant cells also adopt cancer stem cell properties (SOX9). These cells then intravasate through TMEM doorways into the vasculature and become circulating tumor cells (CTCs). **Right Panel:** CTCs retain these programs at the secondary site where the invasion program (Mena^INV^) facilitates extravasation. The dormancy program expressed by these disseminated tumor cells (DTCs) keeps them as single cells.

After extravasation, a tumor cell must survive at a secondary site in order to form metastases. In agreement with other real-time imaging studies (12), we found that a minority of EM injected tumor cells survive in the lung, and only a small fraction of these give rise to micro-metastases within the time frame of our experiments. Though we did not observe a statistically significant difference in the number of cells that eventually progressed to micro-metastases in the EM and SM models, we found that the vast majority of SM cells do survive in the lung as solitary tumor cells, suggesting that they may be in a dormant state. We confirmed this by staining DTCs for NR2F1, a well-established marker for dormancy used in different pre-clinical models (20,21,42,43) as well as in the clinic for breast and prostate cancer patients (41). The co-expression of NR2F1 (a dormancy marker) and SOX9 (a dormancy and cancer stem cell marker) in SM tumor cells suggests that these dormant cancer cells also adopt cancer stem cell properties.

We previously showed that hypoxia in the primary tumor induces NR2F1 expression in a subpopulation of tumor cells (21). Here, we extend this work to find that the primary tumor induces, in disseminating tumor cells, programs of dormancy and stemness (**Figure 6**). Though NR2F1-positive and SOX9^High^ cancer cells constitute a very small percentage of the primary tumor, they become enriched as tumor cells approach TMEM doorways, enter the blood stream, and finally arrest in the lung. Our data indicate that induction of NR2F1, similar to our previous finding for Mena^INV^ (35), is caused by interaction with tumor associated macrophages in a niche surrounding TMEM, and that this programming is carried to the secondary site. This increase in NR2F1-positive cells around TMEM doorways is consistent with previous findings showing that macrophages accumulate in close proximity to the vasculature as tumors progress (68). The increase is also supported by high-resolution intravital imaging studies, which showed that, prior to intravasation, disseminating tumor cells and macrophages rapidly co-migrate (streaming migration) (36) and chemotax towards primary tumor blood vessels along an HGF gradient (69). This mechanism increases the likelihood of interactions between macrophages and tumor cells as they approach TMEM doorways. Taken together, our current work indicates that the NR2F1 expression in tumor cells is induced by contact with macrophages at TMEM doorways.

We have previously shown that depletion of macrophages dramatically reduces the number of tumor cells disseminating to the lung (36, 53). Here, we demonstrated that, within this reduced disseminating population, depletion of macrophages results in a lower level of NR2F1-positivity in both CTCs in the blood, and DTCs in the lung. However, since clodronate treatment depletes macrophages systemically, we cannot definitively rule out a contribution from lung macrophages to conclude that this effect is solely from primary tumor macrophages. Still, two observations indicate that induction of the dormancy program does not occur in the circulation. The first is that a reduction of NR2F1 upon macrophage depletion occurred in both DTCs and CTCs, and the second is that there was no difference between the percentage of NR2F1-positive tumor cells *in vitro* (pre-injection) and in the lungs in the days following injection. Even with this evidence, we cannot rule out that tumor cells could eventually (>3 days) acquire and then maintain a dormancy program at the secondary site via other mechanisms (as we previously demonstrated with TGF-β2 signaling (70)). An important question that remains to be elucidated is the molecular mechanism by which macrophages induce the expression of NR2F1 in tumor cells, but this investigation is beyond the scope of our current study. Finally, we cannot rule out that some programming (e.g. dormancy) may occur via exosomes, as was recently observed in breast cancer metastasis to the bone (71). Nor can we rule out the possibility that the tumor cells we tracked were influenced by the presence of DTCs that had previously arrived in the lung. However, the fact that we observed significant differences in the CTC population indicates that our observations are likely independent of these possible additional influences.

Together, these data indicate that spontaneously disseminating tumor cells acquire programs of dissemination, dormancy, and stemness by interacting with macrophages in the vicinity of TMEM doorways within the primary tumor, and that TMEM doorways are not only sites of tumor cell intravasation (46, 51), but are also microenvironmental niches that program a potentially lethal tumor cell population. This programming imparts to tumor cells the ability to intravasate and extravasate efficiently (via Mena^INV^ expression), to survive long-term, resist anti-proliferative chemotherapy (via dormancy), and to acquire tumor initiation competency (via stemness) that can result in the formation of metastases.

Furthermore, our data reveal a link between dissemination and dormancy. This is an important link because it may explain the observation in patients and in mouse models that early DTCs are dormant and serve as founders of metastasis years after dissemination (72–75).

Finally, the recent identification of TMEM doorways in metastatic sites (24, 76) suggests that the same mechanism of tumor cell programming and dissemination may impact dissemination from secondary on to tertiary sites, even after the removal of the primary tumor. It may therefore be clinically beneficial to inhibit TMEM function systemically even after removal of the primary tumor to prevent this continued systemic spread.

## MATERIALS AND METHODS

### Cell Culture

E0771-GFP medullary breast adenocarcinoma cells, originally isolated from a spontaneous mammary tumor in C57BL/6 mice, were obtained from Dr. Wakefield’s lab at the NIH, who in turn obtained them from Dr. Fengzhi Li in Dr. Enrico Mihich’s lab at Roswell Park Cancer Institute, Buffalo, NY. The MDA-MB-231 human breast cancer cell line was purchased from ATCC. The MDA-MB-231 stably expressing -GFP were generated using retroviral vectors with retroviral packaging, infection and were FACS sorted for the over-expression of each fusion protein, as described elsewhere (77). MDA-MB-231- GFP cells over-expressing Mena^INV^ or Mena11a were generated as previously described (36). The E0771-GFP cell line was cultured in RPMI medium 1640 (ThermoFisher, cat #12633012), media supplemented with 10% (v/v) heat inactivated fetal bovine serum (Atlanta Biologicals, FBS-Premium Select, cat# S11550) and 1% penicillin/streptomycin (Gibco, cat #15-140-122). The MDA-MB-231-GFP, MDA-MB-231-GFP-Mena^INV^ and MDA-MB-231-GFP-Mena11a cell lines were cultured in DMEM (ThermoFisher, cat #12320032) media supplemented with 10% (v/v) FBS and 1% penicillin/streptomycin. Tumor cell lines were used between passage 10-25. The BAC1.2F5 macrophages were cultured in a-MEM supplemented with 10% FBS, and 3000U/mL of purified human recombinant CSF-1 (generously provided by Dr. Richard Stanley, Einstein College of Medicine), and, used at passage 2-15. Human Umbilical Vein Endothelial Cells (HUVECs) were obtained from Lonza and were cultured in EGM-2 SingleQuot Kit media (Lonza, cat #CC-3162) and used at passage 2-10. All cell lines were authenticated at the beginning of the planned experiments, and were routinely tested for mycoplasma and resulted negative (Sigma LookOut Mycoplasma PCR detection kit, cat #MO0035-1KT).

### Animals

All procedures were conducted in accordance with the National Institutes of Health regulation concerning the care and use of experimental animals and with the approval of the with the approval of the Einstein College of Medicine Animal Care and Use Committee (IACUC). Two different animal models were used: an immunocompetent C57BL/6 mouse model and an immunodeficient NUDE mouse model (*Foxn1^nu^/Foxn1^nu^,* Jackson Labs, cat #007850). Two transgenic variants of the C57BL/6 strain of mice were used for intravital imaging: (i) a VeCad-tdTomato mouse expressing the fluorescent protein tdTomato on all endothelial cells, generated by crossing B6.FVB-Tg(Cdh5-cre)7Mlia/J (Jackson Labs, cat#006137) with B6.Cg-Gt(ROSA)26Sortm14(CAG-tdTomato)Hze (Jackson Labs, cat #007914) and (ii) a wild type C57B6 mouse (Jackson Labs, cat #000664). All mice were bred in house. Only female mice between 12 and 24 weeks of age were used for experiments.

### Metastasis Models

#### Experimental Metastasis Model (EM)

E0771-GFP, MDA-MB-231-GFP, MDA-MB-231-GFP- Mena^INV^ or MDA-MB-231-GFP-Mena11a cells were prepared by trypsinizing a 10 cm confluent culture dish of tumor cells and passing them through a 40 μm cell strainer (Falcon, cat #352340) to avoid clumps. A total of 2 × 10^5^ cells was resuspended in 50 μL of sterile PBS and intravenously (iv)-injected via lateral tail vein into a WHRIL-bearing mouse.

#### Spontaneous Metastasis Model (SM)

E0771-GFP or MDA-MB-231-GFP cells were prepared as described above. E0771-GFP cells (1 × 10^6^) were resuspended in 100 μL of sterile PBS, MDA-MB- 231-GFP cells (1 × 10^6^) were resuspended in 100 μL of 20% of collagen I (BD Biosciences, cat #354234). Cells were injected in the 4th lower left mammary fat pad of the mouse. Tumor size was measured once per week using a Vernier caliper and tumor volume was calculated using the ellipsoid formula: tumor volume (mm^3^) = (width in mm)^2^ x (length in mm)/6. When tumor reached a size around 1,500 mm^3^, a WHRIL was placed 24 hours before imaging.

#### Surgery for Implantation of the Window for High Resolution Imaging of the Lung (WHRIL)

The surgery for the WHRIL implantation and the Window passivation method were performed as described previously (24, 78).

### Intravital Imaging

#### Procedure

Mice were anesthetized using 2% isofluorane and injected with 50 µL of 155 kDa Tetramethylrhodamine-labeled dextran (200 µg/mL) retro-orbitally for visualization of blood flow, as previously described(24, 79). Mice were inverted, placed on the microscope stage, and a fixturing plate was taped to the stage using paper tape. The animal was placed in a heated chamber maintained at physiological temperatures by a forced air heater (WPI Inc., AirTherm ATX), during the course of imaging. Imaging was performed on a previously described, custom-built, two-laser multiphoton microscope (80). All images were captured in 16 bit using a 25x 1.05 NA objective lens and acquired with two frame averages.

### Retention of tumor cells analysis

As summarized in **Figure 1A**, for the EM model, the WHRIL was implanted in a naïve mouse and 24-hour post-operation, 2 x 10^5^ E0771-GFP or MDA-MB-231-GFP tumor cells were iv-injected. Tumor cells observed trapped in the lung vasculature under the (WHRIL) were immediately recorded at the time zero (t = 0). Subsequently, every 8 hours the lung was imaged (t = 8, 16, 24, 32, 40, 48, 56, and 64 hours post-injection) and, using the micro-cartography technique to return to the same imaging field(24), the same fields of view were re-localized to observe and track the same tumor cells longitudinally. For the SM model in which E0771-GFP or MDA-MB-231-GFP tumor cells spontaneously disseminated from the primary tumor to the lung, the WHRIL was implanted in a mouse bearing a tumor of *∼*1,500 mm^3^ in size. 24-hours post-operation, the lung was imaged to identify DTCs already present in the lung (**Figure 1A, Time Point = Pre**). These cells were then excluded from further analysis. 8 hours later, the lung was again imaged and any newly arrived DTCs were recorded (**Figure 1A, Time Point = 0 hr**). Then, similar to the EM model, the lung was imaged every 8 hours (at 8, 16, 24, 32, 40, 48, 56, 64 hours from the time that DTCs arrive in the lung vasculature) to track longitudinally the fate of spontaneously DTCs over a period of 64 hours.

For the retention analysis, we defined a tumor cell at each time point as “*retained”* in the lung when we were able to observe the same cell during each imaging session, independent of whether the cell was intra- or extravascular. If the tumor cell was not observed at a time point, then we defined this cell as “*disappeared”*. Kaplan-Meier survival curves showing the retention of tumor cells in the lung over time were generated with the GraphPad Prism software.

### Extravasation of Tumor Cells Analysis

Tumor cells were divided into two subclasses: intravascular or extravascular, based on their location relative to the vasculature. To determine the location of a cell relative to the vasculature, the images of the vasculature at each time point (t = 8, 16, 24, 32, 40, 48, 56, 64 hours) were analyzed and registered with the corresponding prior time point using Adobe Photoshop CC 2015. The tumor cells overlapping with vasculature were considered to be intravascular. Cells not overlapping with the vasculature were considered as having extravasated. Tumor cells were excluded if their localization could not be accurately resolved. For extravascular tumor cells, we were also able to identify three different fates over time: 1) tumor cell death, 2) survival as a single and solitary tumor cell, or 3) formation micro-metastases. We identified tumor cell death when cellular debris (apoptotic bodies) was observed the field of view. We determined a tumor cell to have survived as a single tumor cell when we observed it in the same field of view over time. Finally, we determined a tumor cell to have formed micro-metastases when we observed a cell to become a cluster of more than 5 tumor cells together.

### Migration of Tumor Cells Analysis

To track the migration of tumor cells, and to confirm that we were able to observe the same tumor cell at each imaging session, continuous time lapse imaging of a minimum 8 hours was performed to record the motility path of tumor cells in the lung vasculature. For the time-lapse imaging sessions, a tail-vein catheter was inserted to periodically provide hydration (PBS) and to allow re-administration of dextran (79). Cell motility was manually tracked from one frame to the next using the ROI_Tracker plugin (80). These traces were plotted in Excel (Microsoft) and used to calculate the migration paths of each cell.

### Image Processing and Analysis

Image analysis was performed in ImageJ (81). All images presented are the raw data acquired from the microscope with minimal adjustment of brightness and contrast. Time lapse movies were assembled into Hyperstacks and any slight, residual x-y drift not eliminated by the fixturing window was removed using the HyperStackReg plugin (82) (https://github.com/ved-sharma/HyperStackReg), which is based upon the StackReg plugin for ImageJ (83).

### In Vitro Co-Culture Assay

For the co-culture assay, E0771-GFP tumor cells were plated either in direct contact, or in a 6- well Transwell system with BAC1.2F5 macrophages or HUVEC cells at a 1:4 ratio (20,000 tumor cells and 80,000 macrophages or HUVEC cells) for 24 hours in DMEM supplemented with 10% FBS. The following day, tumor cells were stained for NR2F1 expression as described in the “Immunofluorescence of Tumor Cells Cultured in Vitro” section.

### Immunocytochemistry Staining of Tumor Cells in vitro

To test the baseline expression of NR2F1 or SOX9^High^ in tumor cells cultured *in vitro,* E0771-GFP or MDA-MB-231-GFP cells were plated with a confluence of 70-80% in a 35 mm glass-bottom dish (Ibidi, cat #81156) for 24 hours in DMEM 10% FBS. The following day, tumor cells were washed with PBS three times, fixed in 4% (w/v) paraformaldehyde at room temperature for 15 minutes, permeabilized with 0.15% (v/v) Triton X-100 for 10 minutes, and blocked with a blocking buffer solution (10% FBS, 1% BSA, 0.0025% fish skin gelatin in PBS) at room temperature for 1 hour. Then, tumor cells were incubated overnight at 4 C in the presence of primary antibodies against chicken anti-GFP (Novus, cat #NB100- 1614, concentration 100 μg/mL) and rabbit anti-NR2F1 (Abcam, cat #ab181137, concentration 5 μg/mL), or rabbit anti-SOX9 (Millipore, cat #AB5535, concentration 1 μg/mL). The day after, cells were washed with PBS containing 0.05% Tween-20, and incubated for 1 hour at room temperature with secondary antibodies conjugated with Alexa Fluor 488 for GFP (Invitrogen, cat #A11039, concentration 1 μg/mL) and Alexa Fluor 546 for NR2F1 or for SOX9 (Invitrogen, cat #A11034, concentration 1 μg/mL). Following three washes in PBS containing 0.05% Tween-20, cells were incubated with spectral DAPI (Akoya Biosciences, cat #SKU FP1490), for 5 minutes. Negative controls included incubation with PBS solution instead of the primary antibody. Fluorescence images were captured using an epi-fluorescence microscope (GE, DeltaVision) with a 60x objective, and CoolSNAP HQ2 CCD camera. For image analysis, the NR2F1 channel was thresholded to just above background based upon the negative control. For SOX9^High^ the channel was thresholded so that the number of SOX9^High^ tumor cells was ∼5% of the total number of tumor cells in the primary tumor. The same threshold was applied to the lung tissue analysis.

### Immunofluorescence (IF) Staining of Fixed Tissues

In SM mice, primary tumors and lungs were collected when the tumor reached a size around 1.500 mm^3^. In EM mice, lungs were collected after 3 days post tumor cells injection. After extraction, primary tumors and lungs were fixed in 10 mL of 10% of formalin (v/v) for 48 hours. After 48 hours, tissues were embedded in paraffin and then processed for immunofluorescence staining.

### IF Staining for a single marker: NR2F1, SOX9, or Mena^INV^

Primary tumor or lung paraffin- embedded sections (4 μm) were first melted at 60 C for 1 hour, deparaffinized in xylene, and rehydrated in a graded series of ethanol solutions. Antigen unmasking was performed 1 mM EDTA (pH 8.0) or 1x citrate buffer (pH 6.0) (Diagnostic BioSystems, cat # 99990-096) at 97 C for 20 minutes in a conventional steamer. Slides were rinsed with PBS, permeabilized with 0.5% Triton X-100 in PBS for 5 min at room temperature and incubated with the blocking buffer solution (10% FBS, 1% BSA, 0.0025% fish skin gelatin in PBS) for 1 hour at room temperature. Slides were then incubated overnight at 4 C with primary antibodies against chicken anti-GFP (Novus, cat #NB100-1614, concentration 100 μg/mL) and rabbit anti-NR2F1 (Abcam, cat #ab181137, concentration 5 μg/mL), rabbit anti-SOX9 (Millipore, cat #AB5535, concentration 1 μg/mL), chicken-Mena^INV^ (generated in the Condeelis Laboratory, concentration 0.25 µg/mL), or rat-Endomucin (Santa Cruz, cat #sc-65495, concentration 2 μg/mL). For the Mena^INV^ staining, goat anti-GFP (Novus, cat #NB100-1770, concentration 10 μg/mL) was used. Slides were washed three times in PBS containing 0.05% Tween-20 and incubated with a secondary fluorescent antibody (all Invitrogen, concentration 1 μg/mL) for 1 hour in the dark at room temperature. After washing, slides were incubated with spectral DAPI for 5 minutes and mounted with ProLong Gold antifade reagent (Life Technologies, cat #P36980). Negative controls included incubation with PBS solution instead of the primary antibody. The slides were imaged on the Pannoramic 250 Flash II digital whole slide scanner (3DHistech) using a 20x 0.75NA objective lens. Fluorescence images were captured using an epi- fluorescence microscope (GE, DeltaVision) with a 100x objective, and CoolSNAP HQ2 CCD camera. Total cell numbers per high-power field (65x65 µm^2^, see legend) were counted and the percentages of positive or negative cells were calculated. For image analysis, the NR2F1, Mena^INV^, and Endomucin channels were thresholded to just above background based upon the negative control.

### IF co-staining for double markers: NR2F1 and SOX9, or NF2F1 and IBA1

For the NR2F1 and SOX9, or NR2F1 and IBA-1 co-staining, a multiplex immunofluorescence Perkin Elmer Opal 4-color Fluorescent IHC kit was used according to the manufacturer’s directions. After standard slide preparation as described above, slides were stained with different combinations of primary antibodies. For the NR2F1 and SOX9 co-staining, chicken anti-GFP (Novus, cat #NB100-1614, concentration 10 μg/mL) and rabbit anti-NR2F1 (Abcam, cat #ab181137, concentration 0.5 μg/mL), rabbit anti-SOX9 (Millipore, cat #AB5535, concentration 0.1 μg/mL), were mixed. For IBA-1 staining, rabbit anti-IBA-1 (Wako, cat #019-19741, concentration 0.05 μg/mL) was used. The slides were imaged on the Pannoramic 250 Flash II digital whole slide scanner using a 20x 0.75NA objective lens. Negative controls included incubation with PBS solution instead of the primary antibody. Fluorescence images were captured using an epi-fluorescence microscope (GE, DeltaVision) with a 100x objective and CoolSNAP HQ2 CCD camera. Total cell numbers per high-power field (100×) were counted and the percentages of positive or negative cells were calculated. For image analysis, the NR2F1 and SOX9 were analyzed as above described. IBAI-1 channels were thresholded to just above background based upon intensity. A custom-written ImageJ macro was used to count the number of macrophages in each field of view.

### TMEM Immunohistochemistry Staining

Tumor sections were deparaffinized, as described above, and stained for TMEM. TMEM stain is a triple immunostaining in which 3 antibodies are applied sequentially and developed separately with different chromogens on a Dako Autostainer. TMEM staining was performed as previously described (48). Briefly, we used an anti-pan-Mena antibody (BD, cat. #610693, concertation 5 µg/mL) to detect invasive tumor cells, and anti-IBA1 antibody (Wako, cat. #019-19741, concentration 0.167 µg/mL) to detect macrophages, and an anti-endomucin (Santa Cruz, cat #sc-65495, concentration 0.67 µg/mL) to detect the blood vasculature. TMEM sites in the E0771-primary tumor were identified manually by a pathologist.

### TMEM vs. NR2F1 Positive Tumor Cells Distance Analysis

Sequential sections from primary E0771-GFP tumors were stained for TMEM IHC, as described above, and with IF for both GFP (to detect tumor cells) and NR2F1 (as described in the “Immunofluorescent Staining” section). TMEM IHC and IF images were aligned in ImageJ using the Landmark Correspondences plugin. A series of custom-written ImageJ macros were used to calculate the distance of each NR2F1 positive tumor cell in the field to its nearest TMEM or nearest blood vasculature lacking TMEM. Distance histograms were analyzed and plotted in GraphPad Prism.

### Macrophage Depletion using Clodronate

Tumor bearing mice were treated for 7 days with either 200 µL of PBS liposomes (injected intraperitoneally, or with 200 µL Clodronate liposomes intraperitoneally (Encapsula Nano Sciences, cat. #CLD-8901). Primary tumors or lungs were collected, fixed as described above and paraffin-embedded sections were stained for macrophages or NR2F1 expression as previously described in the “Immunofluorescent Staining - NR2F1 and IBA-1 IF co-staining” section.

### Circulating Tumor Cells Staining

Mice bearing a primary tumor of ∼1000 mm^3^ size were anesthetized with isoflurane and about 1 mL of blood was drawn from the right heart ventricle using 25 G needles coated with heparin. Erythrocytes were lysed using 10 mL of 1x RBC lysis buffer (eBioscience, cat #00-4333-57) for 10 min a room temperature. The samples were centrifuged at 1,000 rpm for 5 min, cells were reconstituted in 10 mL of DMEM supplemented with 10% FBS, plated in a 35 mm glass-bottom dish, and allowed them to adhere overnight. The following day, tumor cells were stained using antibodies against chicken anti-GFP (Novus, cat #NB100-1614, concentration 10 μg/mL) and rabbit anti-NR2F1 (Abcam, cat #ab181137, concentration 5 μg/mL), goat anti-SOX9 (RD, cat #AF3075, concentration 1 µg/mL), as described in the “Immunocytochemistry Staining of Tumor Cells in Vitro” section. Fluorescence images were captured using an epi-fluorescence microscope (GE, DeltaVision) with a 60x objective and CoolSNAP HQ2 CCD camera. For image analysis, the NR2F1 and SOX9^High^ were analyzed as above described.

### Western Blot

Western blot analysis was performed using standard protocols as previously described (84).

### Statistical Analysis

All statistical analysis was carried out using the GraphPad Prism software version 7. Data are expressed as mean ± standard error of the mean (S.E.M). Unless otherwise specified in the figure legends, statistical significances between two groups were determined using unpaired, two-tailed Student’s *t*-tests. Differences were considered significant for *p*<0.05. All *in vivo* and *in vitro* experiments were independently repeated and included at the least three biologically independent samples, as indicated in the legends. For two-group comparisons between experimental and spontaneous metastasis (EM vs SM) models, statistical power calculation was performed using the following parameters: significance level (adjusted for sidedness) of 0.025, total number of mice used in the analysis 6 (i.e. 3 mice per experimental group), and expected difference in means equal to 3 SD units based on the assumption that the SD of the response variable was 1 unit. The probability of type II error in the analyses was calculated to be 0.22 (i.e. statistical power of 78%). For clodronate experiments, statistical power calculation was performed using the following parameters: significance level (adjusted for sidedness) of 0.025, total number of mice used in the analysis 16 (i.e. 8 mice per experimental group), and expected difference in means equal to 1.6 SD units based on the assumption that the SD of the response variable was 1 unit. The probability of type II error in the analyses was calculated to be 0.16 (i.e. statistical power of 84%).

## AUTHOR CONTRIBUTIONS

LB, MHO, JSC & DE designed the experiments, coordinated the project, and prepared figures. LB, JSC, JAA-G, MHO, & DE analyzed and interpreted the data. LB, DE, & AC performed the experiments. VPS wrote the custom ImageJ macro for the distance analyses. GSK and XY helped with statistical analysis. YL performed the TMEM staining. LB, YW, & AC managed the animal colonies. XY designed the levelling plate which enabled the large-volume high-resolution intravital imaging experiments. CLD & XC performed the western blot experiment. JAG provided the protocols for NR2F1 staining. JAA-G & LB analyzed and interpreted the dormancy and metastasis data. ED & DS replicated the data on macrophage induced NR2F1 using alternate methods and provided expertise in NR2F1 detection. LB, JSC, MHO, JAA-G, & DE wrote the manuscript.

The authors would like to thank Peng Guo, Vera DesMarais, and Hillary Guzik for assistance and support in the use of the Perkin Elmer 250 slide scanner and DeltaVision microscope. The authors would like to thank also Dr. Lalage Wakefield’s lab at the NCI for the donation of the E0771 cell line.

This work was supported by the NCI grants: R01-CA218024, S10-OD019961, P30-CA013330, F32- CA243350, T32-CA200561, K12-GM102779, R01-CA109182, R01-CA218024, P30-CA196521, T32-CA078207, The NYS grant DOH01-ROWLEY-2019-00038; grants from METAvivor and HiberCell LLC, The Gruss-Lipper Biophotonics Center and its associated Integrated Imaging Program; and Jane A. and Myles P. Dempsey. JAA-G is a Samuel Waxman Cancer Research Foundation Investigator.

## Competing interests statement

Dr. Julio Aguirre-Ghiso (a Co-author in this article) is a scientific Co-Founder of, Scientific Advisory Board Member, and equity owner in the private company, HiberCell LLC. In addition, Dr. Aguirre-Ghiso receives financial compensation as a consultant for HiberCell LLC. HiberCell LLC. is a Mount Sinai spin-out company focused on the research and development of therapeutics that prevent or delay the recurrence of cancer.

## SUPPLEMENTARY FIGURES & LEGENDS

**Supplementary Figure 1:**
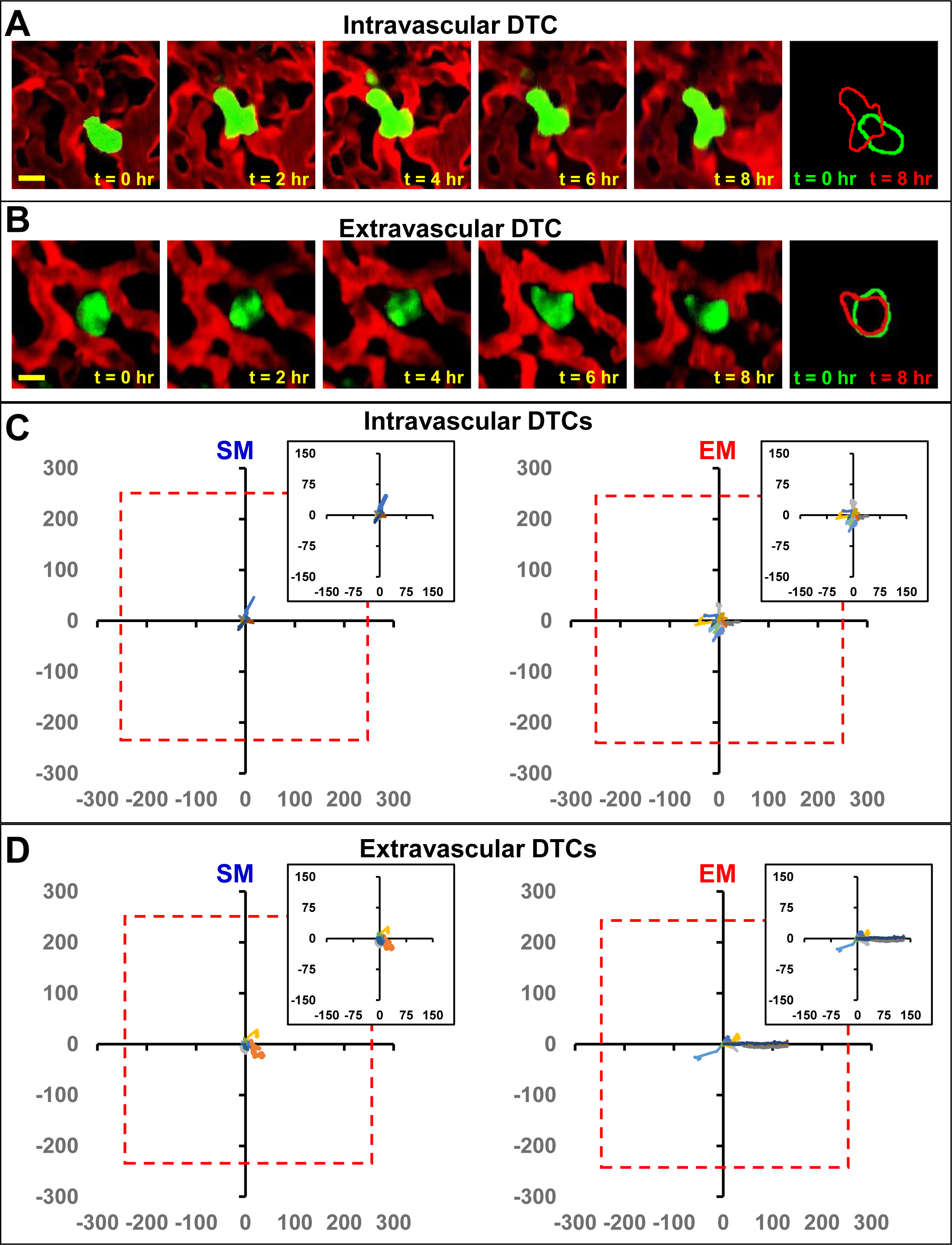
Disseminated Tumor Cells Remain within an Imaging Field of View throughout an 8 hr Period. **A:** Representative intravital microscopy images showing intravascular disseminated tumor cells at different time points spanning 8 hrs. Red = tdTomato labeled endothelial cells and 155 kDa Tetramethylrhodamine dextran labeled blood serum, Green = GFP labeled tumor cells. **Rightmost Panel:** Outlines of the tumor cell at t=0 (green) and t=8 hr (red) showing net displacement. Scale bar = 15 µm. **B:** Representative intravital microscopy images showing extravascular disseminated tumor cells at different time points spanning 8 hrs. Red = tdTomato labeled endothelial cells and 155 kDa Tetramethylrhodamine dextran labeled blood serum, Green = GFP labeled tumor cells. **Rightmost Panel:**Outlines of the tumor cell at t=0 (green) and t=8 hr (red) showing net displacement. **C:** Traces tracking the migration of intravascular E0771-GFP cells within SM (**left**) and EM (**right**) models over an 8 hr period of time. Each tracked tumor cell is represented in a plot with the initial position (t = 0 hrs) translated to the origin so as to provide an overview of the migration path of each cell. Red dashed box indicates a full field of view in the microscope (512 µm. Insets are zoom-ins of the central 150 µm. Average cell velocities: SM = 1.2 ± 0.3 μm/hr; EM = 1.7 ± 0.3 μm/hr; mean ± SEM. SM: n = 11 tumor cells in 4 mice. EM: n = 22 tumor cells in 2 mice. **D:** Traces tracking the migration of extravascular E0771-GFP cells within SM (**left**) and EM (**right**) models over an 8 hr period of time. Each tracked tumor cell is represented in a plot with the initial position (t = 0 hrs) translated to the origin so as to provide an overview of the migration path of each cell. Red dashed box indicates a full field of view in the microscope (512x512 µm^2^). Insets are zoom-in of the central 150 µm. Average cell velocities: SM = 2 ± 0.6 μm/hr; EM =1.8 ± 0.6 μm/hr mean ± SEM. SM: n = 7 tumor cells in 4 mice. EM: n = 11 tumor cells in 3 mice.

**Supplementary Figure 2:**
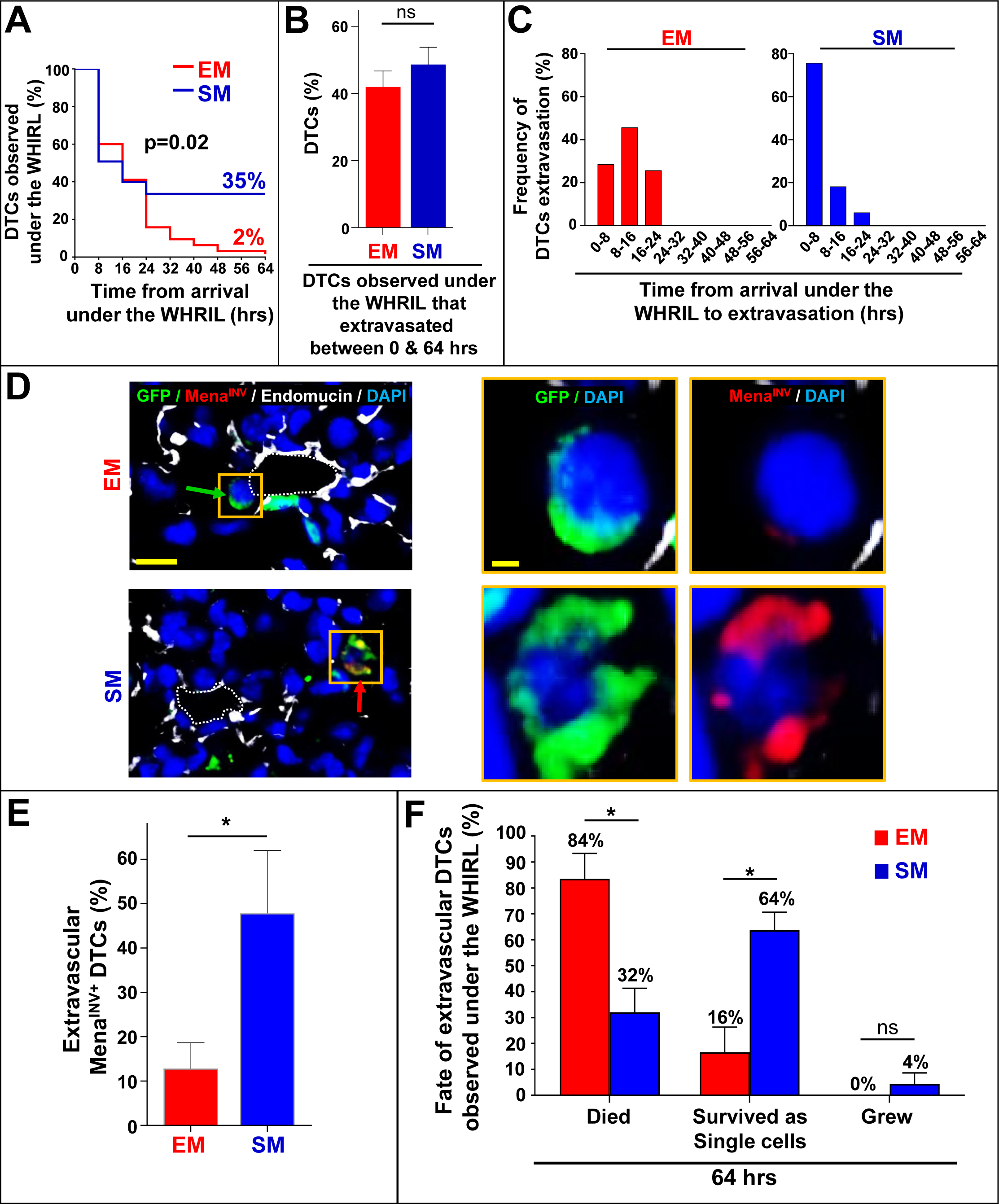
Tumor Cells that Spontaneously Disseminate from the Primary Tumor to the Lung have a Drastically Increased Metastatic Efficiency Compared to Intravenously Injected Tumor Cells. **A:** Kaplan-Meier curves showing the percentage of 231-GFP disseminated tumor cells observed under the WHRIL at each 8 hr time point over a period of 64 hours. EM: n = 92 tumor cells analyzed in 4 mice. SM: n = 67 tumor cells analyzed in 3 mice. **B:** Percentage of 231-GFP EM and SM tumor cells observed under the WHRIL that extravasated between 0 and 64 hrs after arrival. EM: n = 37 tumor cells in 4 mice. SM: n = 33 tumor cells in 3 mice. Bar = mean. Error bars = ±SEM. **C:** Quantification of the time from arrival under the WHRIL to extravasation into the lung parenchyma for each 231-GFP EM and SM tumor cell. **Left:** EM: n = 35 tumor cells in 4 mice. **Right:** SM: n = 33 tumor cells in 3 mice. **D: Left:** Representative immunofluorescence images of Mena^INV^ expression in extravascular 231-GFP tumor cells in the lung of an EM model (**top**) and an SM model (**bottom**). Green arrow: Mena^INV^ negative tumor cell. Red arrow: Mena^INV^ positive tumor cell. Scale bar = 50 μm. **Right:** Zoomed in view of yellow boxed area of a disseminated tumor cell in both models. Green = GFP, Red = Mena^INV^, White = endomucin, Blue = DAPI. Scale bar = 15 μm. **E:** Quantification of extravascular Mena^INV^ positive disseminated tumor cells in the lung of each group from **D**. EM: n = 58 cells in 6 animals. SM: n = 85 cells in 6 animals. Bar = mean. Error bars = ±SEM. * = p<0.05. **F:** Percentage of extravascular 231-GFP disseminated tumor cells that died, survived, or grew after extravasation in EM and SM models 64 hrs (**Left**) and 120 hrs (**Right**) after arrival to the lung vasculature. Bar = mean. Error bars = ±SEM. EM: n = 35 tumor cells in 4 mice. SM: n = 33 tumor cells in 3 mice. * = p<0.05; ns = not significant.

**Supplementary Figure 3:**
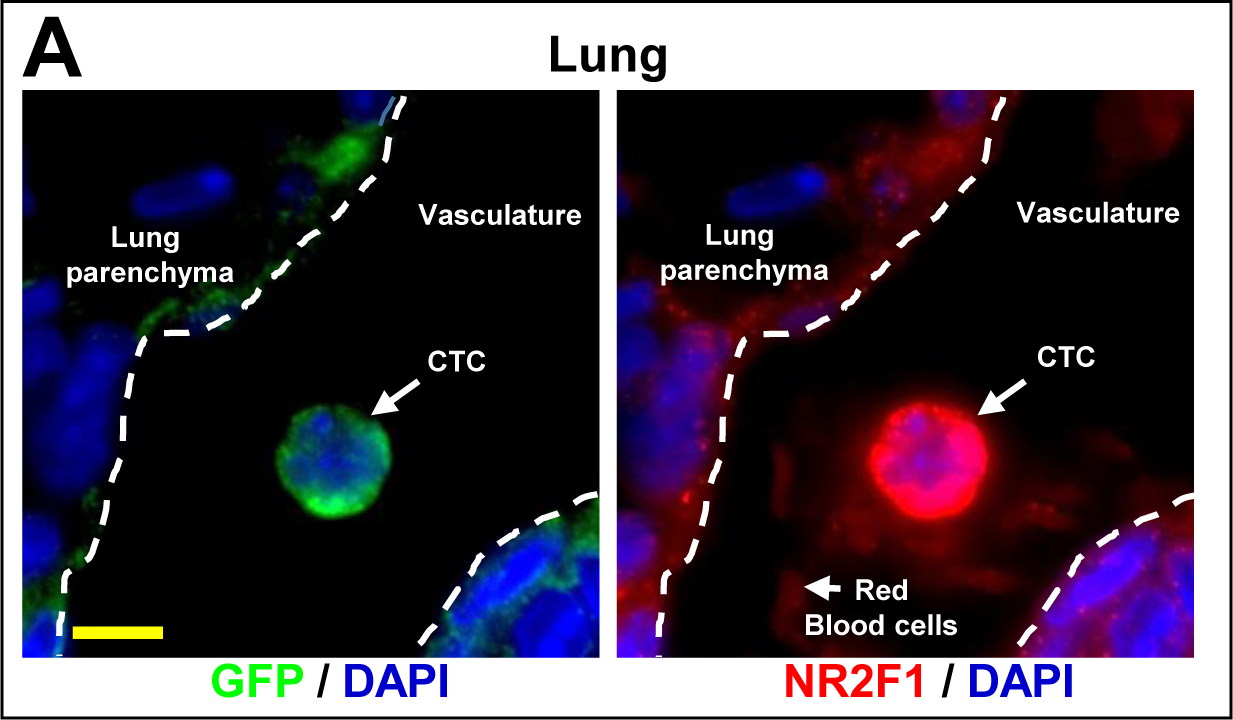
Spontaneously Circulating Tumor Cells Are Positive for NR2F1 When Arriving in the Lung Vasculature. **A:** Representative immunofluorescence image of NR2F1 expression in circulating tumor cells in lung vasculature from a E0771-GFP SM model. **Left:** GFP channel. **Right:** NR2F1 channel. Green = GFP, Red = NR2F1, Blue = DAPI. Dotted line indicates the boundary of the vasculature. CTC = Circulating Tumor Cell. Scale bar = 10 μm.

**Supplementary Figure 4:**
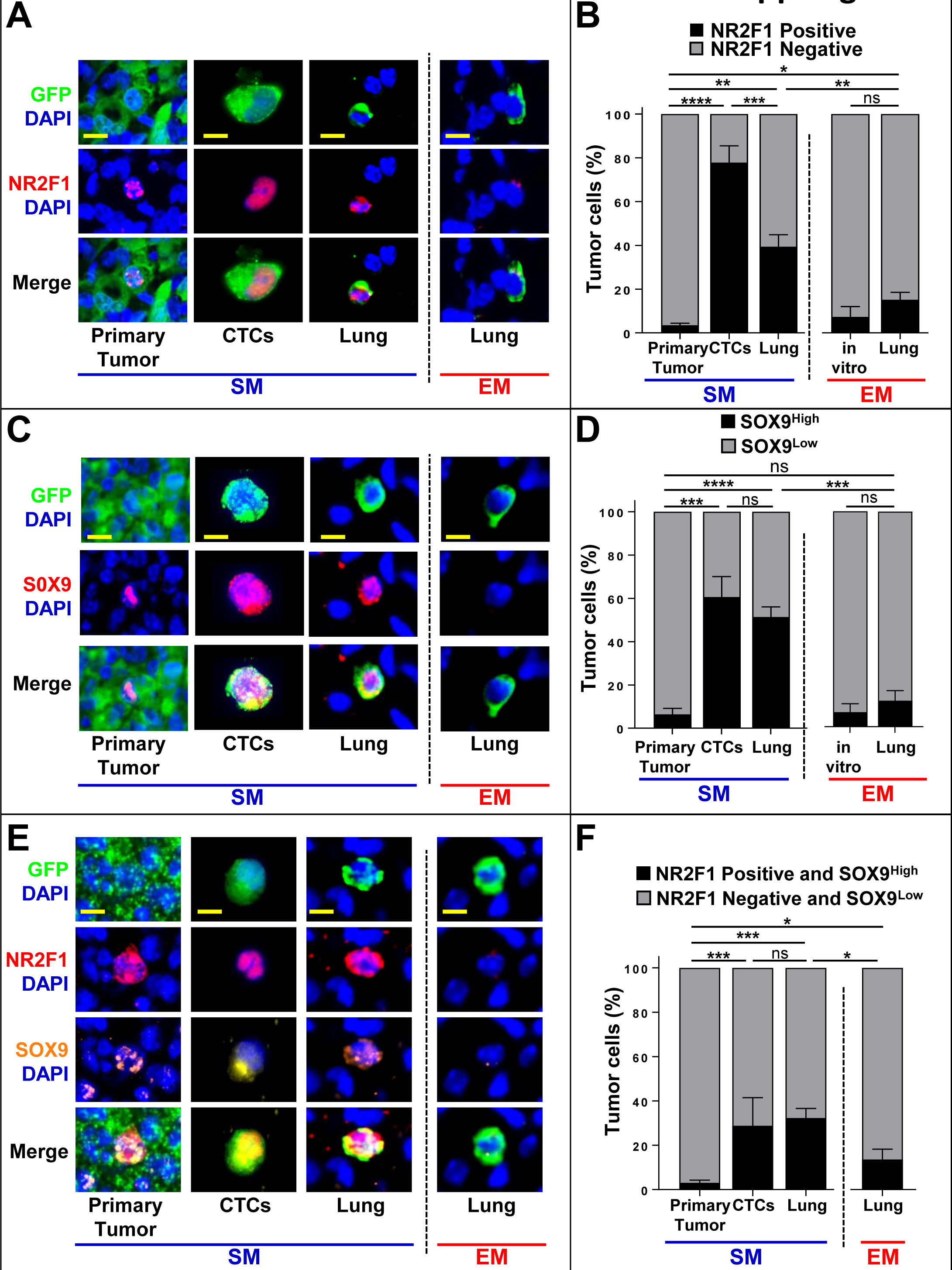
Spontaneously Metastasizing Tumor Cells Are More Frequently Positive for Dormancy and Stem-like Markers Compared to Intravenously Injected Tumor Cells. **A:** Representative immunofluorescence images of NR2F1 expression in primary tumors, circulating tumor cells (CTCs), and disseminated tumor cells (Lung) from a 231-GFP SM model (**Left**) and in disseminated tumor cells (Lung) from an EM model (**right**). Green = GFP, Red = NR2F1, Blue = DAPI. Scale bar for Primary Tumor = 50 μm. Scale bar for CTCs and Lung = 15 μm. **B:** Percentage of NR2F1-positive and negative tumor cells in each group in **A**. Primary Tumor: n = 96 fields of view (65x65 µm^2^) in 6 animals; CTCs: n = 166 cells in 5 animals; SM Lung: n = 113 cells in 6 animals; EM Lung: n = 133 cells in 7 animals. Bar = mean. Error bars = ±SEM. * = p<0.05. ** = p<0.01. *** = p<0.001. **** = p<0.0001. ns = not significant. **C:** Representative immunofluorescence images of SOX9 expression in primary tumors, circulating tumor cells (CTCs), and disseminated tumor cells (Lung) from a 231-GFP SM model (**Left**) and in disseminated tumor cells (Lung) from an EM model (**right**). Green = GFP, Red = SOX9, Blue = DAPI. Scale bar for Primary Tumor = 50 μm. Scale bar for CTCs and Lung = 15 μm. **D:** Percentage of SOX9^High^ tumor cells from each group in **C**. Primary Tumor: n = 100 fields of view (65x65 µm^2^) n = 2643 cells in 5 animals; CTCs: n = 83 cells 2 animals, SM Lung: n = 93 cells in 5 animals; EM Lung: n = 99 cells in 6 animals. In vitro: n = 576 cells in 2 independent experiments. Bar = mean. Error bars = ±SEM. *** = p<0.001. **** = p<0.0001. ns = not significant. **E:** Representative images of triple immunofluorescence staining for GFP, NR2F1, and SOX9^High^ expression in primary tumors, circulating tumor cells (CTCs), and disseminated tumor cells (Lung) from a 231-GFP SM model (**Left**) and in disseminated tumor cells (Lung) from an EM model (**right**). Green = GFP; Red = NR2F1; Orange = SOX9; Blue = DAPI. Scale bar for Primary Tumor = 50 μm. Scale bar for CTCs and Lung = 15 μm. **F:** Percentage of double positive (NR2F1-positive and SOX9^High^) tumor cells from each group in **E**. Primary Tumor: n = 97 fields of view (65x65 µm^2^) in 5 animals; CTCs: n = 118 cells in 5 animals; SM Lung: n = 136 cells in 6 animals; EM Lung: n = 90 cells in 4 animals. Bar = mean. Error bars = ±SEM. * = p<0.05; *** = p<0.001; ns = not significant.

**Supplementary Figure 5:**
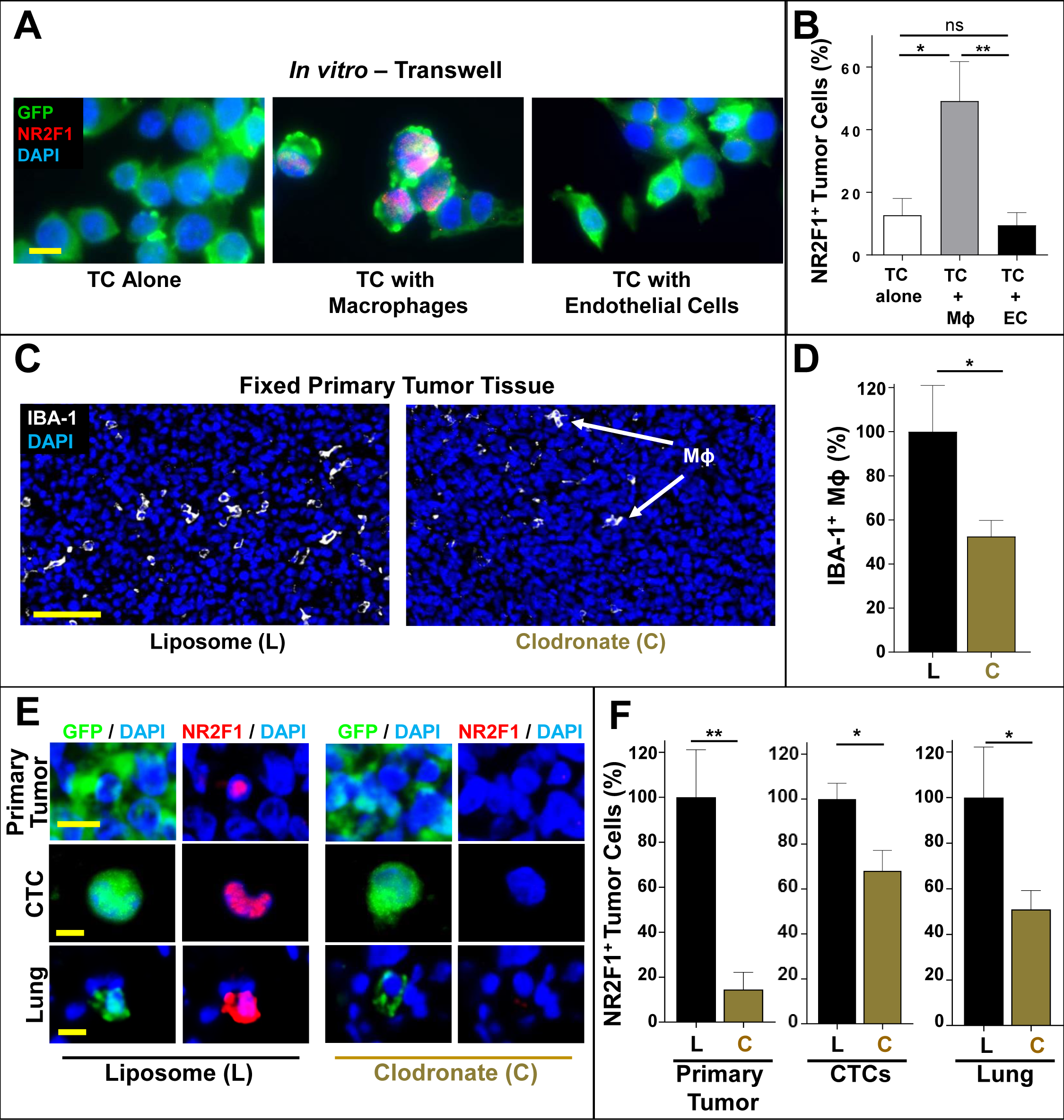
Macrophages Regulate Dormancy in Disseminating Tumor cells. **A:** Representative immunofluorescence images of NR2F1 expression in E0771-GFP tumor cells cultured in a transwell system either alone (**Left**), together with macrophages (**Middle**), or together with endothelial cells (**Right**). GFP = green; NR2F1 = red; DAPI = blue. Scale bar = 15 μm. **B:** Percentage of NR2F1-positive tumor cells from each group in **A**. TC alone: n = 612 cells in 6 independent experiments; TC + Mϕ; n = 680 cells in 5 independent experiments. TC + EC; n = 1370 cells in independent experiments. Bar = mean. Error bars = ±SEM ** = p<0.01; ns = not significant. TC = Tumor Cell. Mϕ = macrophage. EC = Endothelial Cell. **C:** Representative immunofluorescence images of 231-GFP primary tumor tissues treated for 5 days with either control liposomes or clodronate liposomes and stained for macrophages: IBA-1 = White; DAPI = Blue. Scale bar for Tumor = 100 μm. Mϕ = Macrophage. **D:** Number of IBA-1 positive macrophages in 10 fields of view (1102x633 μm) in each group from **A**. Control Liposomes: n = 50 HPFs in 5 animals. Clodronate liposomes: n = 60 HPFs in 6 animals. * = p<0.05. Bar = mean. Error bars = ±SEM. **E:** Representative immunofluorescence images of NR2F1 expression in primary tumors, circulating tumor cells (CTCs), and disseminated tumor cells (Lung) from a 231-GFP SM model treated with control liposomes (L**eft**) or with clodronate liposomes (**Right**). Green = GFP, Red = NR2F1, Blue = DAPI. Scale bar for Primary Tumor = 50 μm. Scale bar for CTCs and Lung = 15 μm. **F:** Percentage of NR2F1-positive tumor cells in each group from **G**. Control Liposomes: Primary Tumor: n = 88 fields of view (65x65 µm^2^) in 5 animals; CTCs: n = 130 cells in 5 animals; Lung: n = 98 cells in 4 animals. Clodronate Liposomes: Primary Tumor: n = 109 fields of view (65x65 µm^2^) in 6 animals; CTCs: n = 337 cells in 6 animals; Lung: n = 126 cells in 6 animals. Bar = mean. Error bars = ±SEM. * = p<0.05. ** = p<0.01. ns = not significant.

**Supplemental Figure 6:**
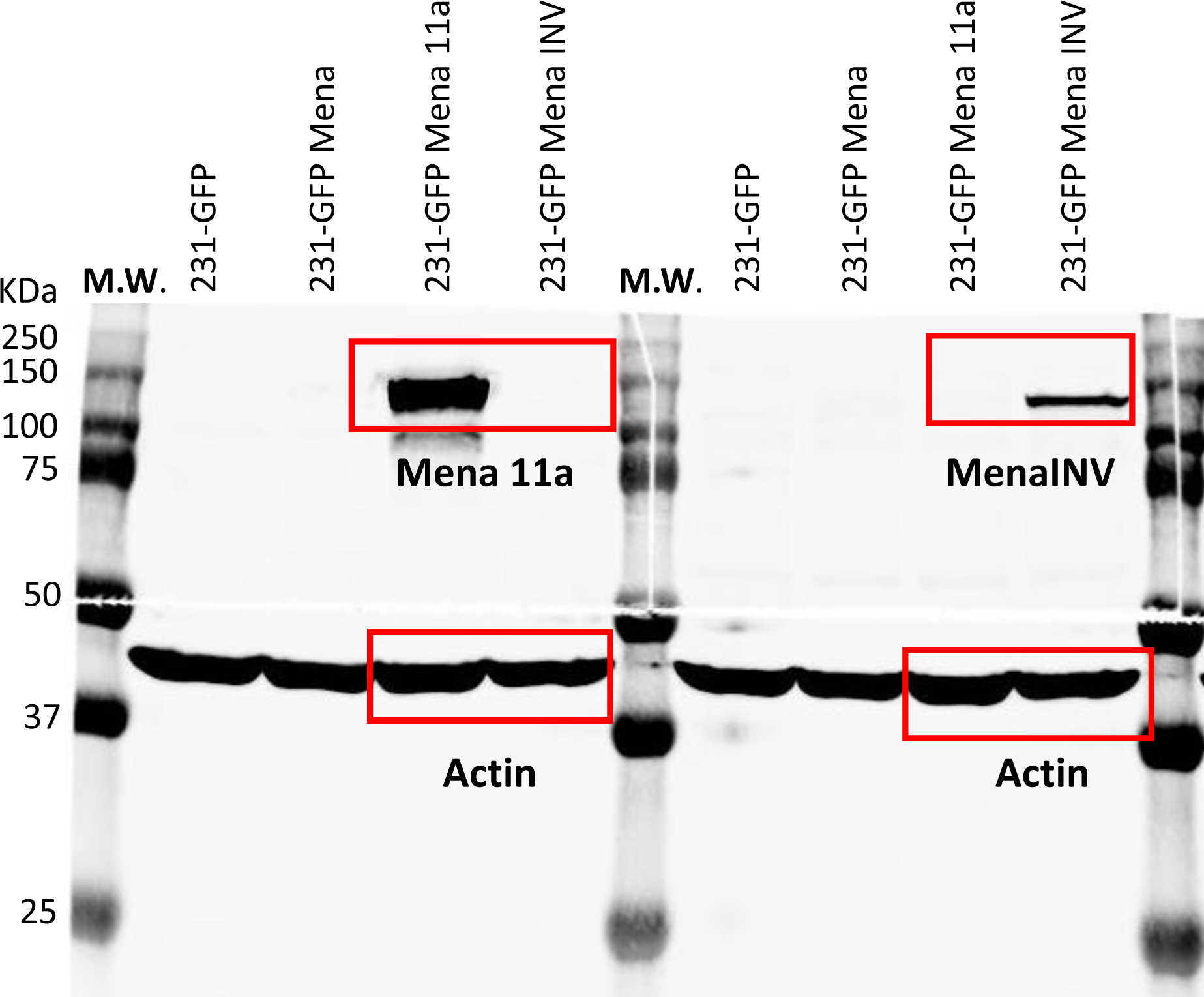
Uncropped blot for Figure 1H.

Movie 1. Time Lapse movie showing the movement of an intravascular disseminated tumor cells. Red (blood) channel was averaged to improve definition of the vascular boundaries. Red = 155kD TMR-dextran labeled vasculature. Green = GFP tumor cell.

Movie 2. Time Lapse movie showing the movement of an extravascular disseminated tumor cells. Red (blood) channel was averaged to improve definition of the vascular boundaries. Red = 155kD TMR-dextran labeled vasculature. Green = GFP tumor cell.

## Notes

https://www.dropbox.com/s/10jksjx2ijjy1hg/Movie%201%20Intravascular%20Tumor%20Cell.avi?dl=0

https://www.dropbox.com/s/w66acc0mxv5ysqu/Movie%202%20Extravascular%20Tumor%20Cell.avi?dl=0

